# Axonal Tau sorting depends on the PRR2 domain and 0N4R-specific interactions hint at distinct roles of Tau isoforms in synaptic plasticity

**DOI:** 10.1101/2024.06.28.601286

**Authors:** M. Bell-Simons, S. Buchholz, J. Klimek, H. Zempel

## Abstract

Tau pathology is a major hallmark of Alzheimer’s disease (AD) and related diseases, called tauopathies. While Tau is normally enriched in axons, somatodendritic missorting of the microtubule-associated protein is a key event in early disease development. Tau missorting promotes synaptic loss and neuronal dysfunction but the mechanisms underlying both normal axonal sorting and pathological missorting remain unclear. Interestingly, the disease-associated Tau brain isoforms show different axodendritic distribution, but the distinct role of these isoforms in health and disease largely unknown. Here, we aimed to identify domains or motifs of Tau and cellular binding partners that are required for efficient axonal Tau sorting, and we studied the differences of the isoform-specific Tau interactome. By using human *MAPT*-KO induced pluripotent stem cell (iPSC)-derived glutamatergic neurons, we analyzed the sorting behavior of more than 20 truncated or phosphorylation-mutant Tau constructs, and we used TurboID-based proximity labelling and proteomics to identify sorting- and isoform-specific Tau interactors. We found that efficient axonal Tau sorting was independent of the N-terminal tail, the C-terminal repeat domains, and the general microtubule affinity of Tau. In contrast, the presence of the proline-rich region 2 (PRR2) was necessary for successful sorting. Our interactome data revealed peroxisomal accumulation of the Tau N-terminal half, while axonal Tau interacted with the PP2A activator HSP110. When we compared the interactome of 0N3R- and 0N4R-Tau, we observed specific interactions of 0N4R-Tau with regulators of presynaptic exocytosis and postsynaptic plasticity, which are partially associated with AD pathogenesis, such as members of the CDC42 pathway and the RAB11 proteins, while 0N3R-Tau bound to MAP4 and other cytoskeletal elements. In sum, our study postulates that axonal Tau sorting relies on the PRR2 domain but not on microtubule affinity, and unravels a potential isoform-specific role in synaptic function and AD-related dysfunction.

## Introduction

The microtubule-associated protein Tau regulates the dynamic assembly and disassembly of microtubule (MT) filaments in human brain neurons (Cleveland et al., 1977b; Gustke et al., 1994; Kadavath et al., 2015; Kanai & Hirokawa, 1995; Weingarten et al., 1975). The six major human Tau isoforms, which originate from alternative splicing of the *MAPT* gene (Cleveland et al., 1977a, 1977b; Goedert et al., 1989), are mainly localized to the axonal compartment of adult neurons (Binder et al., 1985; Kempf et al., 1996). By rapid binding to or detaching from axonal MT filaments depending on its phosphorylation state (Biernat et al., 1993; Bramblett et al., 1993; Drechsel et al., 1992; Janning et al., 2014; Lindwall & Cole, 1984; Yoshida & Ihara, 1993), Tau controls the MT stability and is thus involved in axonal outgrowth, plasticity, cargo transport and other essential neuronal functions (Brandt et al., 1995; Caceres & Kosik, 1990; Dawson et al., 2010; Dixit et al., 2008; Drubin et al., 1985; Ebneth et al., 1998; Esmaeli-Azad et al., 1994; Hirokawa et al., 1988; Kempf et al., 1996; Liu et al., 1999; Samsonov et al., 2004; Takei et al., 2000; Trinczek et al., 1999). Notably, the Tau isoforms vary in their microtubule affinity and are thought to have distinct roles in Tau function and pathology (Bachmann, Bell, et al., 2021; Buchholz & Zempel, 2024; Bullmann et al., 2009).

Tau is associated with a large number of neurodegenerative diseases (Arendt et al., 2016; Guo et al., 2017), which are summarized as tauopathies (Spillantini et al., 1997). In Alzheimer’s disease (AD), the most prevalent tauopathy by far, pathological alterations of Tau are a key disease hallmark (Arendt et al., 2016; Guo et al., 2017; Tracy & Gan, 2018). So-called Tau pre-tangles, high levels of missorted Tau in the somatodendritic compartment with phosphorylation patterns typical for Tau aggregates in AD brains, were detected decades before clinical disease onset (Braak & Del Tredici, 2011; Braak et al., 2011).

Studies in primary rodent neurons revealed that pathological Tau missorting has multiple neurotoxic effects. Axonal transport and function are disrupted (Ishihara et al., 1999; Roy et al., 2005; Zhang et al., 2004), presumably due to microtubule destabilization (Scholz & Mandelkow, 2014) and direct blocking of motor protein-driven transport (Morris et al., 2015; Scholz & Mandelkow, 2014; Tracy & Gan, 2017). Further, dendritic Tau induces postsynaptic microtubule breakdown and spine loss (Lacroix et al., 2010; Zempel et al., 2013; Zempel & Mandelkow, 2014; Zempel & Mandelkow, 2012; Zempel & Mandelkow, 2015).

Given its role in AD pathogenesis, prevention of Tau missorting bears potential for future therapeutic avenues, but more knowledge about the physiological mechanisms of Tau sorting is crucial to understand its failure in disease. To date, several mechanisms are claimed to contribute to axonal Tau sorting in neurons (Zempel & Mandelkow, 2019). There is evidence for anterograde transport of Tau (Scholz & Mandelkow, 2014), either by free diffusion (Konzack et al., 2007), via microtubule-dependent transport (Mercken et al., 1995; Utton et al., 2002; Utton et al., 2005; Zhang et al., 2004) or by potentially unknown interactions. Other reports suggested the possibility of axonal retention of Tau by compartment-specific modifications or interaction partners (Chudobová & Zempel, 2023; Gauthier-Kemper et al., 2018; Hayashi et al., 2012; Hirokawa et al., 1996; Kishi et al., 2005; Zempel et al., 2010; Zhu et al., 2010). Studies with traceable Tau revealed the existence of a Tau-selective retrograde diffusion barrier within the axon initial segment (AIS) (Li et al., 2011; Zempel et al., 2017), the major structure responsible for development and maintenance of neuronal polarity at the proximal axon (Leterrier, 2016, 2018; Rasband, 2010). Notably, this Tau diffusion barrier acts differently than the general AIS-based protein filter since it depends on polymerized microtubules (Li et al., 2011; Van Beuningen et al., 2015; Zempel et al., 2017) but not on filamentous action (f-actin) integrity (Kole et al., 2008; Nakada et al., 2003; Song et al., 2009; Winckler et al., 1999).

The Tau domains and their interaction partners that are responsible for both anterograde or retrograde Tau sorting mechanisms, are largely unknown. Given the reported key role of microtubules in previous studies (Li et al., 2011; Mercken et al., 1995; Scholz & Mandelkow, 2014; Utton et al., 2005; Zhang et al., 2004), the Tau sites of microtubule-binding are obviously suspect. The C-terminal repeat domains directly bind to microtubules (Amos, 2004), but the Tau binding affinity also depends on the flanking domains, especially the proline-rich region 2 (PRR2) (Amos, 2004; Gustke et al., 1994). Generally, phosphorylation of serine or threonine residues of Tau, e.g., at the AT8 motif (S199, S202, and T205) within the PRR2, or at the KXGS motif (S262, S293, S324, S356) within the repeat domains, decreases microtubule affinity of Tau and increases protein mobility (Biernat et al., 1993; Bramblett et al., 1993; Drechsel et al., 1992; Lindwall & Cole, 1984; Yoshida & Ihara, 1993). Hyperphosphorylation of AT8 and KXGS are associated with Tau missorting and aggregation in AD (Bancher et al., 1989; Kaech & Banker, 2006; Kishi et al., 2005; Malia et al., 2016; Mercken et al., 1992; Zempel & Mandelkow, 2014; Zempel et al., 2010).

*In vitro* data from primary rodent neurons showed that recombinant Tau lacking the microtubule-binding repeat domains was still sorted to the axon, despite its high mobility (Iwata et al., 2019). In contrast, exchange of all PRR2 phosphorylation sites by dephosphorylation-mimetic alanine residues prevented anterograde Tau transit (Iwata et al., 2019). This suggests that abnormal somatic dephosphorylation of Tau and thereby tight microtubule binding impairs efficient Tau sorting Studies with KXGS pseudophosphorylation, and the finding of isoform-specific differences in sorting efficiency strengthened the idea that more Tau mobility improves sorting efficiency (Bachmann, Bell, et al., 2021; Zempel et al., 2017). In addition, the N-terminal tail of Tau, which is part of the projection domain of the protein (Amos, 2004; Hirokawa et al., 1988; Kar et al., 2003; Mukhopadhyay & Hoh, 2001), was shown to interact with membrane-bound annexins at the axon and thereby prevents pathological retrograde missorting (Gauthier-Kemper et al., 2018).

However, most previous studies have considerable limitations. They either focused on one direction of Tau trafficking instead of assessing the interplay of sorting mechanisms or offer only fragmentary analyses of selected Tau mutants (Bell et al., 2021; Gauthier-Kemper et al., 2018; Iwata et al., 2019; Li et al., 2011; Zempel et al., 2017). Possible interactors of Tau that are crucial during sorting remain mostly elusive. On the experimental level, most studies used Tau constructs with fused reporter proteins (Gauthier-Kemper et al., 2018; Iwata et al., 2019; Li et al., 2011; Zempel et al., 2017), which can severely impact the intracellular distribution, and non-human neuronal models that fail to resemble key aspects of human Tau physiology (Janke et al., 1999; Kavanagh et al., 2022; Xia et al., 2016).

In our study, we aim to decipher the sorting process of Tau by expressing a library of truncated and modified Tau constructs in human Tau-depleted iPSC-derived glutamatergic neurons. Our inducible lentiviral expression system with equimolar expression of an unbound reporter protein and the absence of endogenous Tau ensures both efficient axonal sorting of overexpressed Tau, an often-faced problem in sorting studies (Xia et al., 2016), and accurate normalization without bias from the reporter protein. In order to reveal interaction partners crucial for successful sorting, we compared the interactome of axonally sorted and non-sorting Tau constructs with the proximity-based TurboID technology. By using this approach, we also examined differences in the interactome of two Tau isoforms since isoform-specific interaction partners are still largely unknown despite the known differences of Tau isoforms in axonal localization and function.

Taken together, our study of factors responsible for axonal Tau sorting and isoform-specific binding partners could help to develop therapeutic approaches, which prevent or slow the Tau-mediated disease progression.

## Methods

### Molecular Biology

#### Generation of lentiviral expression plasmids with Tau^HA^ fragments

Mutant HA-tagged Tau constructs were engineered with site-directed mutagenesis and restriction-based cloning techniques. All fragments were subcloned into the pJET2.1 vector, and transferred to the lentiviral expression vector pUltra-dox with double vector digestion, DNA purification from gel, and ligation. Successful insertion was confirmed by sanger sequencing. All Tau^HA^ constructs were cloned downstream of the cDNA sequence encoding for dTomato, coupled via a P2A peptide sequence.

#### Generation of lentiviral expression plasmids with BirA-Tau^HA^ fusion constructs

HA-tagged BirA-Tau fusion proteins were generated by using the NEBuilder™ HiFi DNA Assembly Kit (NEB). To this end, untagged Tau isoforms and N-terminally HA tagged BirA biotin ligase, derived from a commercially available vector (Addgene, # #107171), were amplified with overlapping ends (∼20 bp) and assembled with the digested lentiviral target vector pUltra-dox. All Tau^HA^ constructs were cloned downstream of the cDNA sequence encoding for dTomato, coupled via a P2A peptide sequence.

### Cell culture

#### Maintenance of human iPSCs

Two different human iPSC lines were used in this study: 1) The WTC11 cell line carrying a doxycycline-inducible murine *Ngn2* transgene (‘WT iPSC-neurons’ (Miyaoka et al., 2014; Wang et al., 2017; Zhang et al., 2013), and 2) a subclone of the above-mentioned line with CRISPR/Cas9-generated biallelic *MAPT* knockout (‘*MAPT*-KO iPSC-neurons’) (Buchholz et al., 2022). Both iPSC lines were cultivated as previously described (Bell et al., 2021; Buchholz, Bell-Simons, Cakmak, et al., 2024). Briefly, cells were cultured on GelTrex™-coated plates (Thermo Fisher) in iPS-Brew XF (StemMACS™, Miltenyi) supplemented with Anti/Anti™ solution (1X, Thermo Fisher). When reaching confluence, the cultures were passaged with Versene™ passaging solution (ThermoFisher), seeded in thiazovivine-supplemented (2 µM, Axon Medchem) iPS-Brew XF, and cultured in thiazovivine-free medium after one day. Cells were grown at 37 °C and 5 % CO2 in a humidified incubator.

#### Differentiation of hiPSC-derived cortical glutamatergic neurons

Differentiation of both iPSC lines into cortical glutamatergic neuronal cultures was performed as previously described (Bachmann, Linde, et al., 2021; Buchholz, Bell-Simons, Cakmak, et al., 2024) with slight adaptations (Fig. S1a). Briefly, cells were seeded in high density (1.5 to 2 x 10^5^ cells per cm³) onto GelTrex-coated plates with doxycycline-containing (2 µg/ml) pre-differentiation medium supplemented with thiazovivine (day −3). After daily medium change without thiazovivine, cells were seeded with lower density (2.5 x 10^4^ cells/cm^3^ for immunostaining, 4 to 5 x 10^4^ cells/cm^3^ for protein harvest) onto culture plates coated with 50 µg/ml Poly-D-Lysine (PDL, Sigma Aldrich) and 20 µg/ml Cultrex™ 3D-laminin (Sigma Aldrich), in doxycycline-containing (2 µg/ml) maturation medium supplemented with 1:100 GelTrex (day 0). Once per week, half of the medium was replaced by fresh GelTrex-free maturation medium. After three weeks, the medium was completely exchanged by doxycycline-free maturation medium (50 % fresh, 50 % conditioned and sterile filtered), followed by weekly changes of half of the medium.

#### Cultivation of HEK293T cells

HEK293T were cultured on uncoated plates in high-glucose DMEM with GlutaMAX (Thermo Fisher) supplemented with 10 % fetal bovine serum (FBS, Biochrom AG) and Anti/Anti (1X). Confluent HEK293T cell cultures were passaged using 0.05 % Trypsin/0.2 % EDTA (Pan Biotech). Cells were grown at 37 °C and 5 % CO2 in a humidified incubator.

#### Lentivirus production

Lentiviral particles were produced as previously described (Buchholz, Bell 2022) with slight modifications. In brief, HEK293T cells were seeded onto uncoated T75 culture flasks. When cultures reached between 60 to 80 % confluency (day 0), medium was changed to 5 ml fresh HEK medium. 10 µg of the lentiviral expression plasmids pUltra (Addgene #24129), pUltra-chili (Addgene #48687), or pUltra-dox (Addgene #58749) were mixed with the helper plasmids pMD2.G (9 µg, Addgene #12259), and psPAX2 (1 µg, Addgene #12260), and 20 µl of polyethylenimine (PEI, Thermo Fisher) in DMEM without supplements. After 20 min incubation, 5 ml fresh HEK medium was added to the cells. After adding 3 ml of fresh HEK medium at day 1, the entire medium was discarded and replaced by 13 ml HEK medium at day 2. For virus harvesting at day 4, the HEK medium was collected, centrifuged for 5 min at 400 g to pellet HEK cell debris, filtered (filter pore size: 0.45 µm), and stored at −80 °C in aliquots. 8 ml of HEK medium were added, and the virus harvesting was repeated at day 5.

For determination of the lentiviral titers, HEK293T cells were seeded onto 24-well plates, transduced after 4-5 hours with different virus dilutions and fixed after 48 h by adding 7.4 % formaldehyde. Cultures for pUltra-dox titering were co-transduced with rtTA-encoding viruses (Addgene # 58750) and supplemented with 2µg/ml doxycycline. After staining with NucBlue™ (ThermoFisher), all cultures were imaged with the ZOE™ fluorescent imaging system (ZOE, Leica), and the proportion of dTomato-positive neurons was determined with Fiji/ImageJ (V1.53t, NIH). Only cultures with proportions between 5 and 30 % positive neurons were used to avoid biased measurements. The titer was calculated as follows:

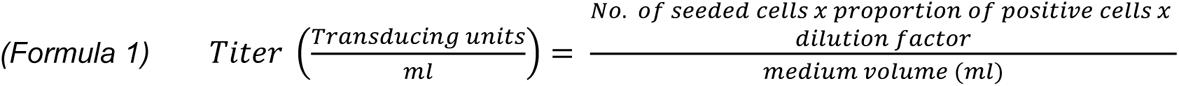

#### Transduction of WT iPSC-neurons

WT iPSC-neurons were transduced with lentiviruses containing empty pUltra or pUltra-chili vectors for constitutive GFP or dTomato expression at different ages as previously described (Buchholz, Bell-Simons, Cakmak, et al., 2024). The virus particles were thawed on ice and mixed with pre-warmed and pH-adjusted maturation medium (50 % fresh, 50 % conditioned and sterile filtered). After cultures were washed once with DMEM/F12 medium, the virus mix was added for 16-24 h and replaced with maturation medium (50/50), containing 2 µg/ml doxycycline when cultures were younger than 3 weeks, otherwise without doxycycline. Young neurons (week 1 to 3) were transduced 3 to 4 days, older neurons (week 4 to 10) 7 to 10 days before fixation.

#### Tau^HA^ construct expression in *MAPT*-KO iPSC-neurons

6-weeks-old *MAPT*-KO iPSC-neurons were transduced with lentiviruses containing pUltra-dox vectors with Tau^HA^ constructs as previously described (Buchholz, Bell-Simons, Cakmak, et al., 2024) In brief, titrated virus particles were thawed on ice and mixed with pre-warmed and pH-adjusted maturation medium (50 % fresh, 50 % conditioned and sterile filtered). After cultures were washed once with DMEM/F12 medium, the virus mix was added for 16-24 h and replaced with doxycycline-containing (0.5 µg/ml) maturation medium (50/50). For protein isolation, 4,000 TU were applied to one 10 cm culture dish with 3 x 10^6^ cells. For immunofluorescence, 100 to 200 TU were applied to cultures in 24-well plates with coverslips. Half of the medium was replaced after seven days. Cells were fixed or harvested 12-13 days after transduction stop.

#### BirA-Tau^HA^ construct expression and biotinylation in transduced KO iPSC-neurons

Transduction and expression of BirA-Tau^HA^ was performed as described above for Tau^HA^ constructs, with modifications. Virus particles were administered in biotin-free maturation medium, B27 was replaced by NS21 supplement (Pan Biotech) and DMEM/F-12 by Neurobasal. For protein isolation, 10,000 TU were applied to one 10 cm culture dish with 3 x 10^6^ cells. For immunofluorescence, the same density (340 TU/well) was applied to cultures grown in 24-well plates with coverslips. After 12-13 days, the medium was exchanged by fresh maturation medium containing 500 µM biotin (Sigma-Aldrich), and neurons were incubated for 20 minutes. Control samples were incubated with biotin-free medium. After incubation, neurons were immediately fixed or harvested.

### Proteinbiochemistry

#### Protein isolation

The protein isolation was adapted from previous studies (Cho et al., 2020) with slight modifications as follows. *MAPT*-KO iPSC-neurons were washed twice with cold DBPS, and lysed by adding ice-cold RIPA buffer (Thermo Fisher) supplemented with 1X cOmplete protease inhibitor cocktail (Sigma-Aldrich) and 1 mM phenylmethylsulfonyl fluoride (PMSF, Sigma-Aldrich) for 1.5 min. The pre-lysed cells were then scraped, collected in low-binding centrifuge tubes (Thermo Fisher), homogenized by multiple pipetting, and lysed for another 20 minutes shaking at 4 °C. The lysates were centrifuged for 10 minutes at 13,000 g, and supernatant was stored at −80 °C. Protein concentration was determined using a bichinonic acid (BCA) protein assay kit (Thermo Fisher) according to the manufacturer’s protocol.

#### Western blot analysis

Isolated proteins were diluted in 6x protein loading buffer containing 0.33 M Tris-HCl (pH 6.8), 34 % glycerol, 10 % SDS, 0.09 % DTT and 0.12 % bromophenol blue (Cho et al., 2020) and boiled for 10 minutes at 95 °C. Proteins were separated using a 10 % polyacrylamide running gel (Mini-Protean system, BioRad), and blotted to PVDF membranes (Bio-Rad) by wet transfer at 30 V overnight at 4 °C. The membrane was blocked in 5 % non-fat dry milk, incubated with the primary antibody (mouse anti-HA (16B12) (1:5000, #901533, BioLegend)) in 5 % milk or 3 % BSA overnight at 4 °C, washed thoroughly, and incubated with the corresponding HRP-coupled secondary antibody (Thermo Fisher) for 1-2 hours at RT. Chemiluminescent signals were generated with the SuperSignal West Pico solutions (Thermo Fisher) and detected using a ChemiDoc XRS+ system (Bio-Rad). Protein levels were measured with HRP-coupled anti-GAPDH (ab9385, Abcam) and anti-vinculin (#18799, Cell Signaling) antibodies, which were incubated for 1 hour at RT.

### Imaging Analysis

#### Fixation & Immunostaining

Neuron cultures were fixed and stained as previously described (Buchholz, Bell-Simons, Haag, & Zempel, 2024). In brief, 7.4 % formaldehyde (FA) were added for 30 minutes, and fixed cultures were long-term stored in 60 % glycerol at −20 °C. For immunostaining, the cultures were permeabilized and blocked with 5 % bovine serum albumin (BSA, Sigma-Aldrich) and 0.1 % Triton X-100 (AppliChem) for 5-10 minutes, incubated with the primary antibodies either for 2-3 h at 20-22 °C or at 4 °C for 16-18 h. The following primary antibodies were used in this study: rabbit anti-Tau (K9JA) (1:1000, #A0024, DAKO), chicken anti-MAP2 (1:2000, ab5392, Abcam), mouse anti-HA (16B12) (1:1000, #901533, BioLegend). After thorough washing steps, the cultures were incubated with the corresponding secondary antibody coupled to AlexaFluor™ fluorophores (Thermo Fisher) for 1-2 h at 20-22 °C. For biotin detection, a NeutrAvidin-AF647 conjugate was generated as described previously (Cho et al., 2020). Nuclei were stained with NucBlue™ (1 drop/ml, Hoechst 33342, TFS) for 20-30 minutes, and cultures were mounted with aqueous PolyMount™ (#18606, Polysciences) onto microscope slides. After 24 h of drying, slides were stored at 4 °C in the dark.

#### Microscopy

Neuron cultures were imaged with the ZOE fluorescent imaging system (Bio-Rad) during differentiation. Images of immunostained neurons were taken on the widefield fluorescence microscope Axioscope 5 (Zeiss), connected to an Axiocam 503 mono (Zeiss) and Colibri 7 LED (Zeiss). Exposure time and light intensity were optimized to prevent sample bleaching and saturation of fluorescent signals. For experiments with Tau^HA^ fragments and BirA-Tau^HA^ fusion constructs, the exposure and light settings were kept identical for all neurons from the same replicate to enable correlation analysis of expression levels and axonal sorting efficiency. Further, exposure times for dTomato and Tau^HA^ or BirA-Tau^HA^ were always set to the same value to allow for the calculation of relative expression ratios. The Zen Blue pro software was used for image acquisition, and Fiji/ImageJ (V1.53t, NIH) was used for quantitative analysis.

#### Sorting analysis of endogenous Tau/MAP2

For axonal sorting analysis of endogenous Tau and MAP2, all images were blinded using the Fiji blind analysis tool. Next, an arbitrary proportion of the soma without nuclear overlap was manually set as the somatic region of interest (ROI). The axonal ROI comprised a segment of 5-10 µm length at 75-100 µm distance to the soma, the segment for the dendritic ROI was 5-10 µm long in 25-50 µm distance to the soma. The nuclear ROI was set by encircling large proportions of the NucBlue-positive area. For all ROIs, the mean fluorescence intensity of anti-Tau, anti-MAP2 and the reporter protein dTomato/GFP was measured using an automated measurement macro (see Suppl. Script 2). Background ROIs in close proximity to the four ROIs were measured and subtracted from the raw ROI values. To obtain the axonal enrichment factor (AEF) of Tau and MAP2, first the ratio of axonal ROI and somatic ROI was calculated for Tau and MAP2, and then divided by the axon-to-soma ratio of dTomato. For the dendritic (DEF) and nuclear enrichment factor (NEF), the respective ROIs were used instead of the axonal ROIs. Neurons were excluded from analysis when they did not reach the signal intensity thresholds set for all four ROIs in all three channels. As an exception, neurons from week 1 and 2 were included in AEF and NEF analysis even with sub-threshold signals of dendritic ROIs. Finally, all neurons were unblinded, and the AEF, DEF, and NEF values were averaged per replicate by using the following formula:

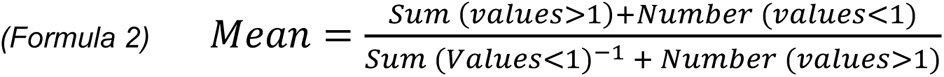

#### Sorting analysis of Tau^HA^ and BirA-Tau^HA^ constructs

The sorting analysis of Tau^HA^ constructs was done like for endogenous Tau with slight differences. To determine unspecific binding of the anti-HA antibody, signals of three randomly chosen non-transduced neurons of the same sample were measured, averaged and subtracted from all ROIs prior to ratio calculation. Neurons were excluded when not reaching thresholds set for all four ROIs (MAP2 & dTomato cannels), or when the signals were not larger than two standard deviations of the unspecific signals (Tau^HA^ channel).

The sorting analysis of BirA-Tau^HA^ constructs was done like for Tau^HA^ constructs with slight differences. Signals from unspecific binding were determined and subtracted for anti-HA and the NeutrAvidin-AF647 staining. Neurons were excluded when not reaching thresholds set for all four ROIs (dTomato cannel), or when the signals were not larger than two standard deviations of the unspecific signals (Tau^HA^ & NeutrAvidin channel).

#### Relative protein levels of Tau^HA^ and BirA-Tau^HA^ constructs

Tau^HA^ and BirA-Tau^HA^ constructs were co-expressed with the 2A-coupled dTomato in equimolar ratios. For calculating the relative protein levels of our constructs after 12-13 days of expression, somatic and axonal ROIs of the anti-HA channel were summed up for each neuron and divided by the somatic/axonal ROI sum of dTomato in the same neuron. The protein levels of each replicate were averaged by using formula 1.

#### Correlation of Tau^HA^/BirA-Tau^HA^ expression levels and the AEF

The correlation between the protein levels of the Tau^HA^ and BirA-Tau^HA^ constructs and the axonal sorting efficiency was examined with simple linear regression analysis. To this end, the AEF values and protein levels were replicate-wise normalized to the maximum AEF and protein level to account for inter-replicate variation, and all neurons were grouped and analyzed together for each construct.

### Interactome analysis

#### Sample preparation

The sample preparation was adapted from previous studies (Cho et al., 2020) with slight modifications. 200 µg protein lysate of each sample were incubated with streptavidin-coated magnetic beads (Thermo Fisher) overnight under rotation at 4 °C. After multiple washing steps, 2 M urea in 50 mM Tris-HCl (pH 7.5) with 1 mM dithiothreitol (DTT, Thermo Fisher) and 5 µg/ml trypsin (Thermo Fisher) were added to the eluate for on-bead digestion for 1 hour shaking at 25 °C. Disulfid bonds were reduced with 4mM DTT for 40 minutes shaking at 25 °C, and alkylated with 10 mM chloroacetamide (CAA) for 50 minutes shaking at 25 °C in the dark. In-solution digestion was performed by adding 0.5 µg trypsin for overnight incubation shaking at 25 °C. Enzymatic activity was stopped by adding formic acid to a final concentration of 1 %. For sample purification, the eluate was centrifuged for 5 minutes at 13,000 g, and the supernatant was loaded onto equilibrated SDB-RP C18 StageTips (Thermo Fisher). After loading the peptides, StageTips were washed, centrifuged at full speed to completely dry, and stored at 4 °C for subsequent LC-MS analysis.

#### LC-MS/MS analysis

All samples were analyzed by the CECAD proteomics facility (CECAD institute, Cologne) on a Q Exactive Plus Orbitrap mass spectrometer that was coupled to an EASY nLC (both Thermo Scientific). Peptides were loaded with solvent A (0.1% formic acid in water) onto an in-house packed analytical column (50 cm length, 75 µm inner diameter, filled with 2.7 µm Poroshell EC120 C18, Agilent). Peptides were chromatographically separated at a constant flow rate of 250 nL/min using the following gradient: initial 3 % solvent B (0.1% formic acid in 80 % acetonitrile), 3-5% B within 1.0 min, 5-30% solvent B within 65.0 min, 30-50% solvent B within 13.0 min, 50-95% solvent B within 1.0 min, followed by washing and column equilibration. The mass spectrometer was operated in data-dependent acquisition mode. The MS1 survey scan was acquired from 300-1750 m/z at a resolution of 70,000 and 20 ms maximum injection time. The top 10 most abundant peptides were isolated within a 1.8 Th window and subjected to HCD fragmentation at a normalized collision energy of 27%. The AGC target was set to 5e5 charges, allowing a maximum injection time of 110 ms. Product ions were detected in the Orbitrap at a resolution of 35,000. Precursors were dynamically excluded for 10.0 s.

All mass spectrometric raw data were processed with MaxQuant (version 2.2.0.0, (Tyanova, Temu, & Cox, 2016)) using default parameters against the Uniprot canonical human database (UP5640, downloaded 05.08.2020) with the match-between-runs option enabled between replicates. Sample normalization was performed as previously described (Cox et al., 2014). Follow-up analysis was done in Perseus 1.6.15 (Tyanova, Temu, et al., 2016). Protein groups were filtered for potential contaminants. Remaining IDs were filtered for data completeness (100 % valid values in at least one group) and missing values imputed by sigma downshift (0.3 σ width, 1.8 σ downshift). Next, all sample groups were filtered for IDs that were enriched (log_10_(p-value)>1.3, log_2_ (fold change)>0.58) compared to both control groups, onlyBirA^HA^ and only0N3R^HA^. The unfiltered list of all detected proteins, and the protein lists after background filtering are available (Tables S1a,b). For the remaining IDs, FDR-controlled two-sided t-tests were performed between all sample groups.

Gene ontology (GO) term analyses was performed to characterize proteins that were enriched in BirA-0N3R^HA^ vs. BirA-Nterm^HA^ or for BirA-0N4R^HA^ vs. BirA-Nterm^HA^. Further, GO term analysis of the combined interactome of BirA-0N3R^HA^ and BirA-0N4R^HA^ was performed to validate found Tau interaction partners. All GO term analyses were done with gProfiler (http://biit.cs.ut.ee/gprofiler/gost). Detailed outcomes of all gProfiler queries are available (Tables S4-S8). Graphical illustration using volcano plots for interactome analysis and dot blots for GO terms analysis were conducted with the ggplot2 package (Version: 3.5.1) in RStudio (Version: 2024.04.1) (see Suppl. Script 1) (Team, 2023; Wickham H & Müller K, 2019).

## Results

### Endogenous-like axonal sorting of exogenous 0N3R-Tau^HA^ in human *MAPT*-KO iPSC-neurons

Poor axonal targeting of exogenous Tau is an often-faced problem in primary rodent cultures and may impede accurate Tau sorting research (Iwata et al., 2019; Xia et al., 2016). In order to obtain a suitable neuronal model for our Tau sorting study, we differentiated human WTC11 iPSCs with a doxycycline-inducible *Ngn2* cassette (Miyaoka et al., 2014; Zhang et al., 2013) with and without biallelic *MAPT* knockout (Buchholz et al., 2022) (Fig. S1a) into cortical glutamatergic neurons using protocols adapted from previous publications (Buchholz, Bell-Simons, Cakmak, et al., 2024; Tracy et al., 2022; Wang et al., 2017)).

First, we assessed the sorting efficiency of endogenous Tau in neurons without *MAPT*-KO (‘WT iPSC-neurons’) at different stages of differentiation. For this, we transduced WT iPSC-neurons with lentiviruses harboring empty pUltra or pUltra-chili vectors for constitutive GFP or dTomato expression and fixed at week 1, 2, 3, 4, 6 and 10 for immunofluorescence analysis (Fig. 1a). The axonal enrichment of Tau, measured as the ratio of endogenous Tau and volume marker (GFP, dTomato) in the axon compared to soma (AEF, see methods for details) increases strongly after week 1, stabilizes by week 6, and elevates towards week 10 with high replicate variability, while MAP2 shows no axonal enrichment at any age (Fig. 1b). The timing of efficient Tau sorting coincides with formation of the axon initial segment (AIS), which occurs around day 14 in human iPSC-neurons (Lindhout et al., 2020)(unpublished data). Both Tau and MAP2, showed modest dendritic enrichment (DEF) compared to the soma with MAP2 being more enriched only at week 3 (Fig. S1b). The considerable dendritic Tau enrichment could be caused by maturation deficits or by technical difficulties of single-dendrite measurements in dense neuronal cultures. Both Tau and MAP2 showed weak nuclear localization without detectable changes during differentiation but the nuclear enrichment (NEF) of MAP2 was slightly higher, (Fig. S1c).

**Fig. 1:**
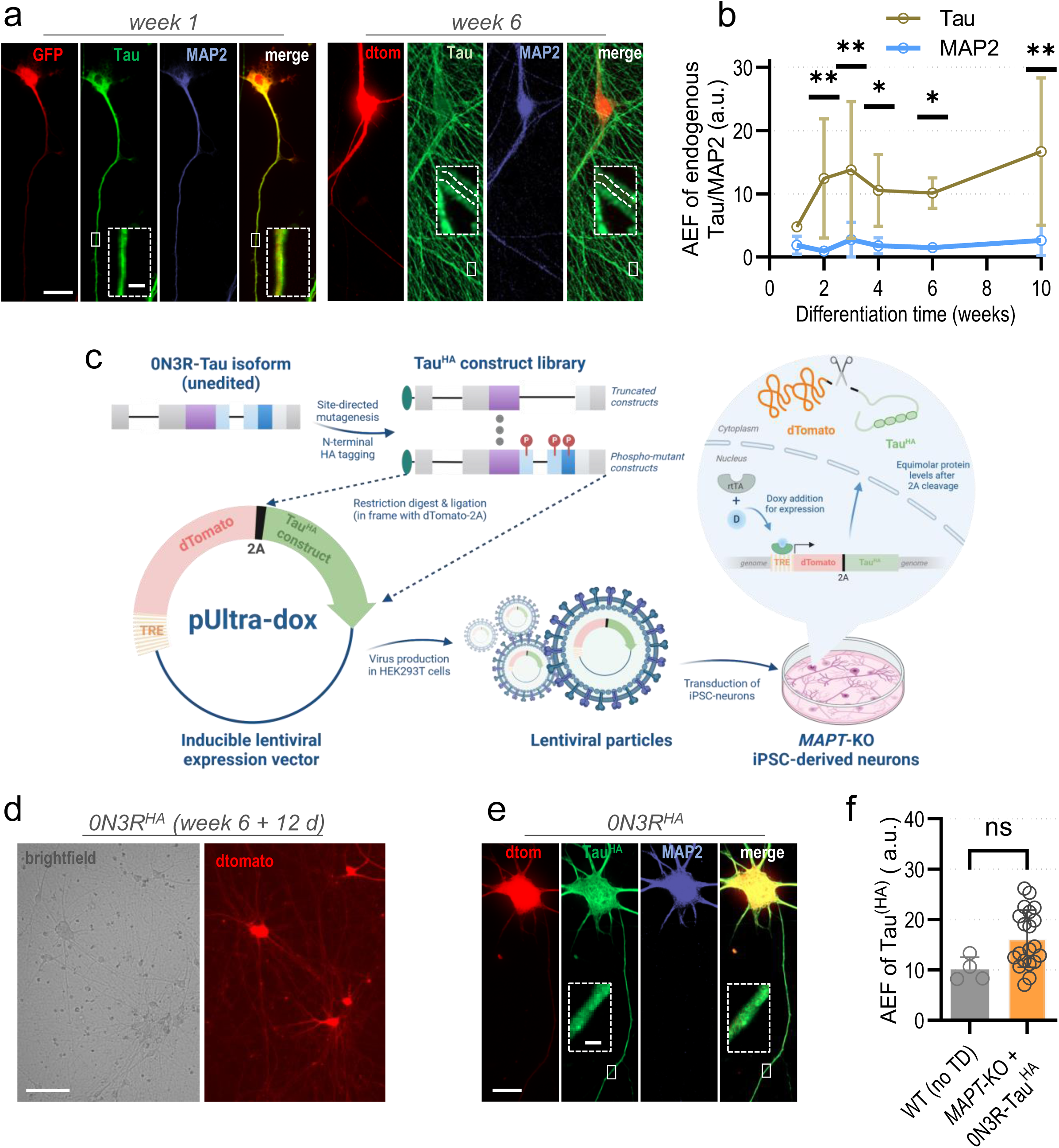
Differentiation of human iPSC-derived neurons and lentivirus production. (a) Human Ngn2-transgenic WTC11 iPSCs were differentiated into iPSC-neurons (see methods) and transduced with lentiviruses that harbour the pUltra or pUltra-chili vector for constitutive GFP or dTomato expression, respectively, at different time points. Cultures were fixed at week 1, 2, 3, 4, 6 and 10, and immunostained. The signals for Tau (green), MAP2 (blue) and dTomato/GFP (red) are shown in single channels and merged (green+red). Axonal sections are magnified (dashed boxes) in merged images. Dashed lines inside the magnifications 6-week-old neurons indicate the axon. Scale bar: 20 µm, scale bar within dashed box: 2 µm. (b) Axonal enrichment (AEF) of endogenous Tau and MAP2 during differentiation normalized to dTomato. Quantification was done for three to five independent experiments (coloured dots) with 8-21 cells per experiment. The coloured dots indicate the arithmetic mean, the error bars show SD. An ordinary two-way ANOVA with Tukey’s correction for multiple comparisons was performed to determine significance levels between different weeks and between two AEF values of the same week. Significance levels: * p < 0.05, ** p < 0.01, ns: p ≥ 0.05. (c) Workflow of Tau^HA^ library sorting analysis. The recombinant Tau^HA^ library was generated with site-directed mutagenesis, and Tau^HA^ constructs were cloned into the lentiviral transfer vector pUltra-dox in frame with the 2A-coupled reporter dTomato under regulation of a tetracycline response element (TRE). Virus particles were produced and harvested from HEK293T cells and transduced to differentiated iPSC-neurons (see Fig. S1a) carrying a biallelic *MAPT*-KO, resulting in undirected genomic integration of the expression cassette. Addition of doxycycline (doxy) 24 h after transduction leads to binding of the reverse tetracycline-activated transactivator (rtTA) to the TRE and equimolar expression of Tau^HA^ constructs and dTomato. (d) *MAPT*-KO iPSC-neurons 12 days after transduction with an empty pUltra-dox vector and right before harvesting (week 6 + 12 days). The upper row shows brightfield, the lower row dTomato signals. Scale bar: 100 µm. (e) *MAPT*-KO iPSC-neurons after expression of 0N3R-Tau^HA^ and dTomato for 12 days and subsequent immunostaining. The signals for Tau^HA^ (green), MAP2 (blue), and dTomato (red) are shown in single channels and merged (green+red). Axonal sections are magnified (dashed boxes) in merged images. Scale bar: 20 µm, scale bar within dashed box: 2 µm. (f) Axonal enrichment (AEF) of 0N3R-Tau^HA^ in *MAPT*-KO iPSC-neurons compared to AEF of endogenous Tau in in 6-weeks-old WT iPSC-neurons. Quantification for was done for four (WT) or 20 (*MAPT*-KO) independent experiments (black dots) with 9-21 cells per experiment. The coloured bars indicate the arithmetic means, the error bars show SD. An unpaired t-test was performed to determine significance levels between both groups. Significance levels: ns: p ≥ 0.05.

Next, we checked the axonal sorting efficiency of HA-tagged 0N3R in Tau-depleted *MAPT*-KO iPSC-derived neurons (“*MAPT*-KO iPSC-neurons”. For this, we cloned 0N3R^HA^ into the doxycycline-inducible lentiviral expression vector pUltra-dox allowing for tunable protein expression, coupled to the reporter protein dTomato via a P2A peptide (Fig. 1c). The produced lentiviral particles were transduced into 6-weeks-old *MAPT*-KO iPSC-neurons with low numbers to ensure infection of single viruses, and doxycycline was added to induce equimolar expression of Tau^HA^ constructs and dTomato for 12-13 days (Fig. 1c,d). We quantified axonal sorting of 0N3R^HA^ (Fig. 1e) and compared it to that of endogenous Tau (Fig. 1f). Strikingly, 0N3R^HA^ sorted with high efficiency, similar or even more efficient than endogenous Tau. The dendritic enrichment of 0N3R^HA^ was not altered and the nuclear levels were still low but modestly increased (Fig. S2b,c). MAP2 showed more axonal enrichment after 0N3R^HA^ overexpression (Fig. S2d), and higher dendritic levels compared to the WT iPSC-neurons (Fig. S2e) but not to the *MAPT*-KO iPSC-neurons, while nuclear levels were not changed (Fig. S2f). The correlation analysis between sorting and protein levels revealed a modest positive correlation, indicating slightly better axonal targeting for higher protein amounts (Fig. S2g), which was rather unexpected.

Taken together, we achieved endogenous-like sorting of exogenous 0N3R^HA^ in *MAPT*-KO iPSC-neurons by using an inducible lentiviral expression system. We now applied our workflow for sorting analysis to our library of truncated and modified Tau^HA^ constructs that we generated with site-directed mutagenesis (Fig. 1c).

### Mutant Tau^HA^ constructs show robust expression levels and highly efficient 2A cleavage

Before we measured the intracellular sorting behavior, we validated the correct expression of all mutant Tau^HA^ constructs with two different approaches. We checked for presence of erroneously uncleaved dTomato-Tau^HA^ fusion proteins and for possible degradation of our mutant Tau^HA^ species post expression by Western blot analysis, and for protein levels relative to dTomato on the single cell level by immunostainings.

In the Western blot, the control 0N3R^HA^ appeared at the expected size of ∼55 kDa, a larger band of uncleaved dTomato-2PA-0N3R-Tau^HA^ was barely visible at ∼95 kDa, suggesting efficient cleavage of the P2A (Fig. 2b, Fig. S3a-c, Fig. S6e). For most of the truncated Tau^HA^ constructs (Fig. 2a), cleaved bands at the expected sizes were visible. For Nterm^HA^, only3R^HA^, and only4R^HA^, no distinct bands were detectable (Fig. 2b). These constructs might be truncated or degraded due to internal protein quality control. For only3R^HA^ and only4R^HA^, this is supported by the decrease in relative protein levels compared to 0N3R-Tau^HA^ (Fig. S3d). The missing band of Nterm^HA^ appears elusive since the protein levels determined by individual neuron analysis are only slightly reduced (Fig. S3d). Interestingly, we saw three very close but distinct bands for noPRR2-Tau^HA^, possible due to variations in phosphorylation or other posttranslational modifications (PTM) that might explain distinct protein sizes. Apart from only3R^HA^ and only4R^HA^, the protein levels were also reduced for the N-terminus-lacking Cterm^HA^ and PRR-3R^HA^, which comprises the microtubule-binding domain. Protein levels for the other constructs were not different from 0N3R^HA^ levels (Fig. S3d). The correlation analysis between protein levels and axonal sorting showed no connection, except for Cterm^HA^, PRR+Cterm^HA^ and only4R^HA^, all positively correlated, i.e. showing increased sorting for higher protein amounts (Fig. S3e,f).

**Fig. 2:**
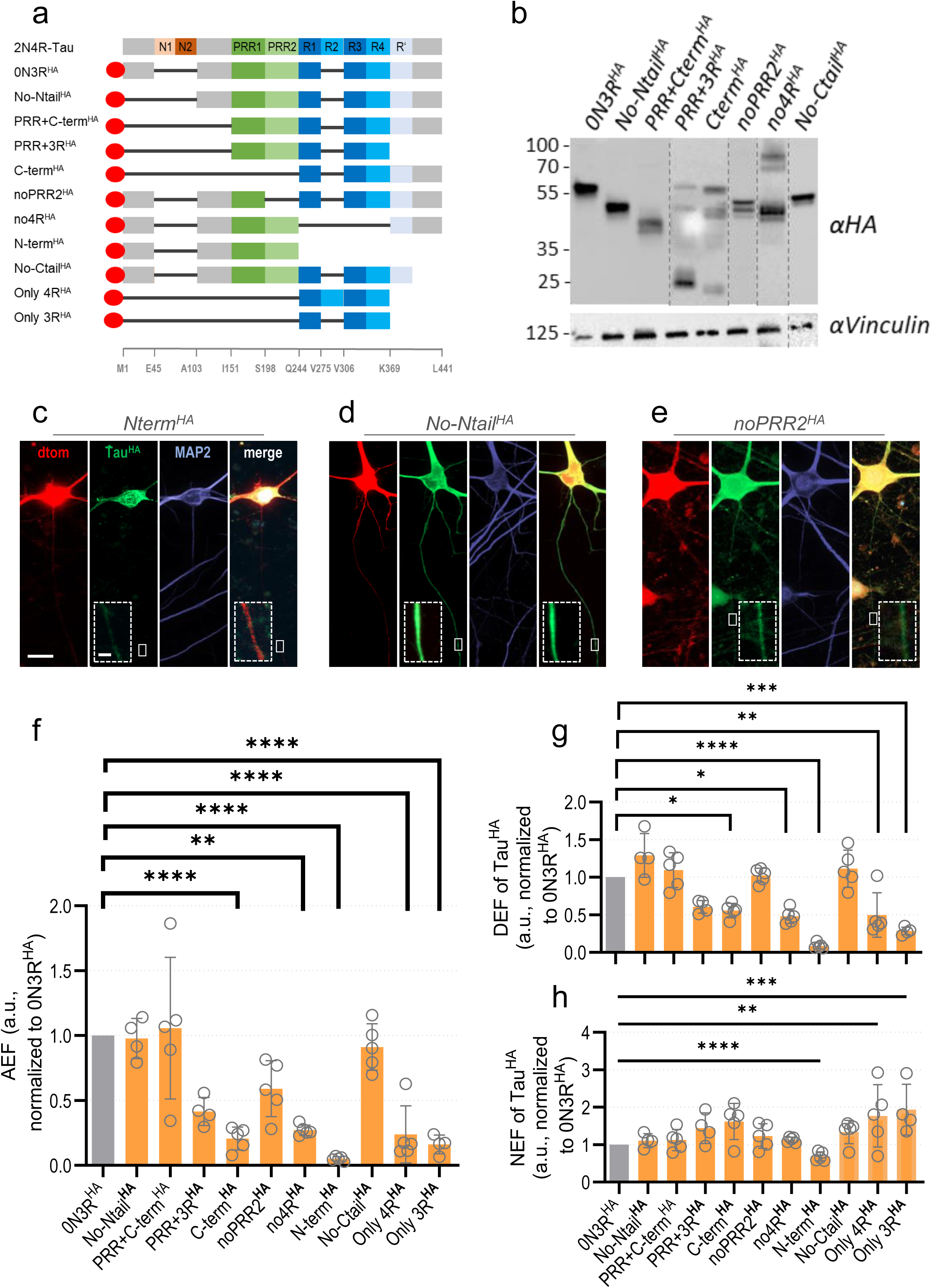
Axonal sorting of transduced 0N3R-Tau^HA^ and truncated Tau constructs. (a) All Tau^HA^ constructs with domain truncation. The 2N4R-Tau isoform (top) and selected amino acid positions (bottom) are given for orientation (see also Fig. S2a). Black bars symbolize the lack of the respective domain, the red oval indicates the HA tag. Brief construct names are indicated at the left. Ntail/Ctail = N/C-terminal tail, Nterm/Cterm = N/C-terminal half, PRR = proline-rich region. (b) Western blot of *MAPT*-KO iPSC-neurons that were transduced with differently truncated Tau^HA^ constructs and harvested after 12-13 days of expression. Vinculin was used as loading control. Protein expression was confirmed for all constructs except for N-term^HA^, only3R^HA^ and only4R^HA^. Notably, noPRR2^HA^ exhibits two very close but distinct bands. The relative abundance of uncleaved dTomato fusion products (see Fig. S4c) compared to cleaved constructs is negligibly low (see Fig. S2a for quantification). Lane contrast was partially adjusted for visualization (dotted lines indicate crop sites, see Fig. S4c-e for raw blots). Note that western blot signals were not used for determining protein level due to different transduction efficiencies. (c-e) *MAPT*-KO iPSC-neurons after expression of Nterm^HA^ (c), No-Ntail^HA^ (d), or noPRR2^HA^ (e) and dTomato for 12-13 days and subsequent immunostaining. The signals for Tau^HA^ (green), MAP2 (blue) and dTomato (red) are shown in single channels and merged (green+red). Axonal sections are magnified (dashed boxes) in merged images. Scale bar: 20 µm, scale bar within dashed box: 2 µm. (f) Axonal enrichment (AEF) of truncated Tau^HA^ constructs in KO iPSC-neurons after 12-13 days expression, normalized to AEF of 0N3R-Tau^HA^. (g) Dendritic enrichment (DEF) of truncated Tau^HA^ constructs in KO iPSC-neurons after 12-13 days expression, normalized to DEF of 0N3R-Tau^HA^. (h) Nuclear enrichment (NEF) of truncated Tau^HA^ constructs in KO iPSC-neurons after 12-13 days expression, normalized to NEF of 0N3R-Tau^HA^. Quantification for (f-h) was done for four to five independent experiments (black dots) with 8-19 cells per experiment. The bars indicate the arithmetic mean, error bars show SD. A mixed-effects model with Dunnett’s correction for multiple comparisons was performed to determine significance levels between 0N3R-Tau^HA^ and all Tau^HA^ constructs. Non-normalized ratios of 0N3R-Tau^HA^ and truncated Tau^HA^ constructs were used for the statistical analysis. Significance levels: * p < 0.05, ** p < 0.01, *** p < 0.001, **** p < 0.0001, ns: p ≥ 0.05.

Neuronal cultures expressing AT8-mutant Tau^HA^ (Fig. 3a) constructs all exhibited distinct bands at the expected size of 0N3R-Tau^HA^. Higher molecular bands of uncleaved fusion protein were negligibly faint (∼1-3 %), except for no4R^HA^ (∼10 % uncleaved) suggesting again the almost complete P2A cleavage (Fig. 3b, Fig S4a,b Fig. S6e). The relative protein levels of all AT8-mutant Tau^HA^ constructs appeared not changed compared to 0N3R-Tau^HA^ (Fig. S4c). Correlation analysis showed positive correlation of protein levels and axonal sorting for S199E^HA^, T205E^HA^, AT8-allE^HA^, and AT8-allA^HA^. Negative correlation was not found (Fig. S4d,e).

**Fig. 3:**
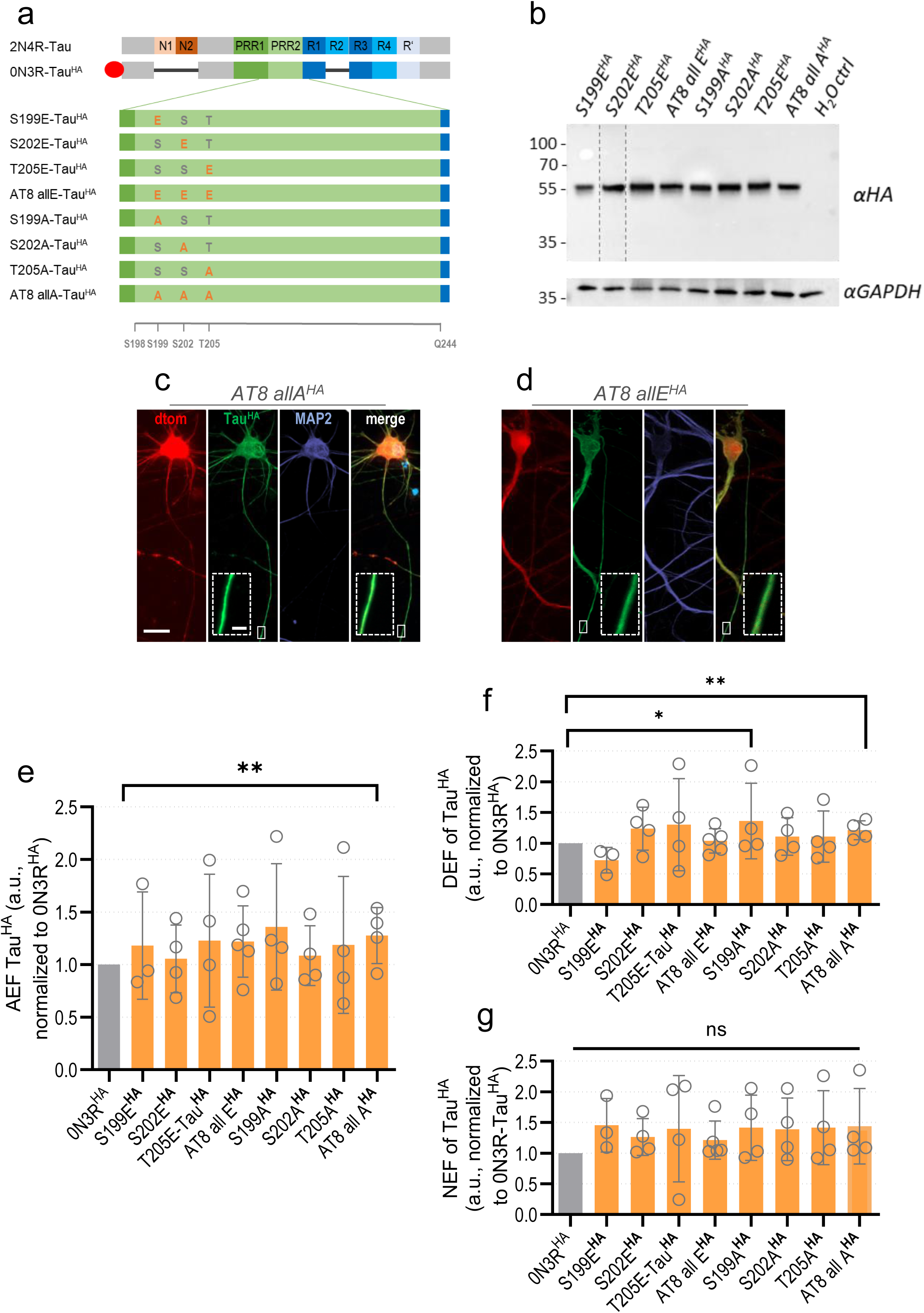
Axonal sorting of transduced 0N3R-Tau^HA^ and AT8-mutant Tau^HA^ constructs. (a) All Tau^HA^ constructs with one or multiple pseudo-phosphorylated (Ser/Thr to Glu) or phosphorylation-deficient (Ser/Thr to Ala) residues within the AT8 motif. The 2N4R-Tau isoform (top) and selected amino acid positions (bottom) are given for orientation (see also Fig. S2a). Black bars symbolize the lack of the respective domain, the red oval indicates the HA tag. Brief construct names are indicated at the left. S = Serine, T = Threonine, A = Alanine, E = Glutamic acid. (b) Western blot of *MAPT*-KO iPSC-neurons transduced with AT8-mutant Tau^HA^ constructs. GAPDH was used as loading control. All bands of AT8-mutant Tau^HA^ constructs appear at the expected size of 55 kDa. The relative abundance of uncleaved dTomato fusion products (see Fig. S6c) compared to cleaved constructs is negligibly low (Fig. S2a for quantification). Lane contrast was partically adjusted for visualization (dotted lines indicate crop sites, see Fig. S6c for raw blots). Note that western blot signals were not used for determining protein level due to different transduction efficiencies. (c-d) *MAPT*-KO iPSC-neurons after expression of AT allA^HA^ (c) or AT8 allE^HA^ (d) and dTomato for 12-13 days and subsequent immunostaining. The signals for Tau^HA^ (green), MAP2 (blue) and dTomato (red) are shown in single channels and merged (green+red). Axonal sections are magnified (dashed boxes) in merged images. Scale bar: 20 µm, scale bar within dashed box: 2 µm. (e) Axonal enrichment (AEF) of AT8-mutant Tau^HA^ constructs in KO iPSC-neurons after 12-13 days expression, normalized to AEF of 0N3R-Tau^HA^. (f) Dendritic enrichment (DEF) of AT8-mutant Tau^HA^ constructs in KO iPSC-neurons after 12-13 days expression, normalized to DEF of 0N3R-Tau^HA^. (g) Nuclear enrichment (NEF) of AT8-mutant Tau^HA^ constructs in KO iPSC-neurons after 12-13 days expression, normalized to NEF of 0N3R-Tau^HA^. Quantification for (f-h) was done for four to five independent experiments (black dots) with 8-19 cells per experiment. The bars indicate the arithmetic mean, error bars show SD. A mixed-effects model with Dunnett’s correction for multiple comparisons was performed to determine significance levels between 0N3R-Tau^HA^ and all Tau^HA^ constructs. Non-normalized ratios of 0N3R-Tau^HA^ and AT8-mutant Tau^HA^ constructs were used for the statistical analysis. Significance levels: * p < 0.05, ** p < 0.01, *** p < 0.001, **** p < 0.0001, ns: p ≥ 0.05.

In neurons with KXGS- and double-mutant Tau^HA^ constructs (Fig. 4a), we could consistently see distinct double bands around 50 kDa with roughly equal proportions (3xKXGA^HA^, 3xKXGE^HA^, AT8+KXGS all E^HA^, AT8+KXGS all A^HA^) or with a dominant larger band (1xKXGA^HA^, 1xKXGE^HA^) (Fig. 4b, Fig. S5a-c). Higher molecular bands indicating uncleaved fusion proteins were almost not detectable for any construct (Fig. 4b, Fig. S5a-c, Fig. S6e). The relative protein levels of all KXGS- and double-mutant Tau^HA^ constructs appeared not changed compared to 0N3R-Tau^HA^ in immunostaining analysis (Fig. S5d).

**Fig. 4:**
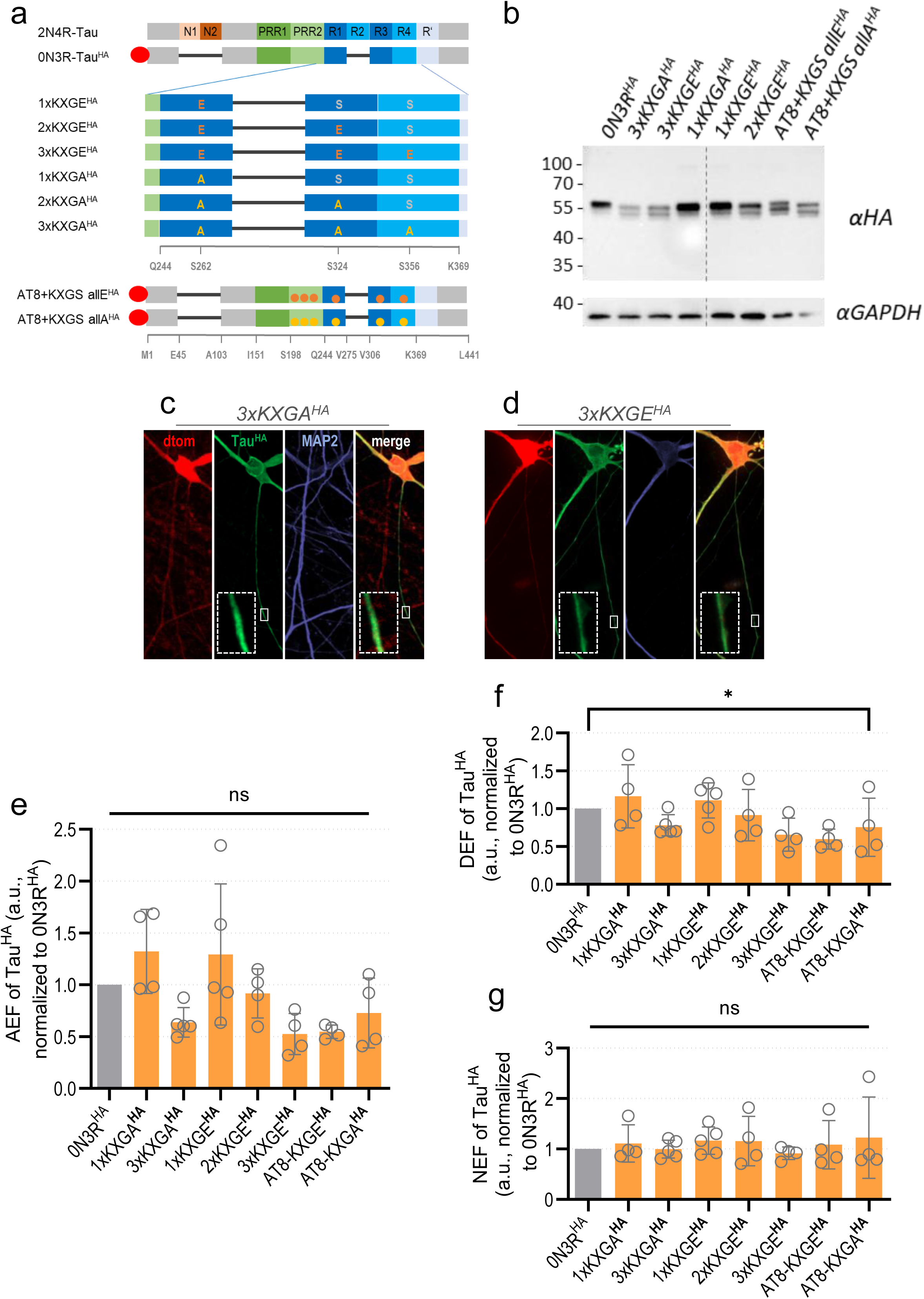
Axonal sorting of transduced 0N3R-Tau^HA^ and KXGS- and double-mutant Tau^HA^ constructs. (a) All Tau^HA^ constructs with one or multiple pseudo-phosphorylated (Ser/Thr to Glu) or phosphorylation-deficient (Ser/Thr to Ala) residues within the KXGS motif, and double KXGS-AT8 mutants. The 2N4R-Tau isoform (top) and selected amino acid positions (bottom) are given for orientation (see also Fig. S2a). Black bars symbolize the lack of the respective domain, the red oval indicates the HA tag. Brief construct names are indicated at the left. S = Serine, T = Threonine, A = Alanine, E = Glutamic acid. (b) Western blot of *MAPT*-KO iPSC-neurons transduced with KXGS- and double-mutant Tau^HA^ constructs. GAPDH was used as loading control. All bands of AT8-mutant Tau^HA^ constructs appear at the expected size of 55 kDa. Remarkably, all constructs showed two very close but distinct bands of various intensity ratios, possibly due to differential posttranslational modification. The relative abundance of uncleaved dTomato fusion products (see Fig. S6c) compared to cleaved constructs is negligibly low (Fig. S2a for quantification). Lane contrast was partically adjusted for visualization (dotted lines indicate crop sites, see Fig. S6c for raw blots). Note that western blot signals were not used for determining protein level due to different transduction efficiencies. (c-d) *MAPT*-KO iPSC-neurons after expression of 3xKXGA^HA^ (c) or 3xKXGE^HA^ (d) and dTomato for 12-13 days and subsequent immunostaining. The signals for Tau^HA^ (green), MAP2 (blue) and dTomato (red) are shown in single channels and merged (green+red). Axonal sections are magnified (dashed boxes) in merged images. Scale bar: 20 µm, scale bar within dashed box: 2 µm. (e) Axonal enrichment (AEF) of KXGS- and double-mutant Tau^HA^ constructs in KO iPSC-neurons after 12-13 days expression, normalized to AEF of 0N3R-Tau^HA^. (f) Dendritic enrichment (DEF) of KXGS- and double-mutant Tau^HA^ constructs in KO iPSC-neurons after 12-13 days expression, normalized to DEF of 0N3R-Tau^HA^. (g) Nuclear enrichment (NEF) of KXGS- and double-mutant Tau^HA^ constructs in KO iPSC-neurons after 12-13 days expression, normalized to NEF of 0N3R-Tau^HA^. Quantification for (f-h) was done for four to five independent experiments (black dots) with 8-19 cells per experiment. The bars indicate the arithmetic mean, error bars show SD. A mixed-effects model with Dunnett’s correction for multiple comparisons was performed to determine significance levels between 0N3R-Tau^HA^ and all Tau^HA^ constructs. Non-normalized ratios of 0N3R-Tau^HA^ and KXGS- and double-mutant Tau^HA^ constructs were used for the statistical analysis. Significance levels: * p < 0.05, ** p < 0.01, *** p < 0.001, **** p < 0.0001, ns: p ≥ 0.05.

In brief, the investigated Tau^HA^ constructs were correctly expressed in our *MAPT*-KO iPSC-neurons without signs of protein degradation. Yet, lower protein levels were observed for the truncated Tau^HA^ constructs, including Cterm^HA^, Nterm^HA^, and PRR+3R^HA^, while only3R^HA^ and only4R^HA^ were not detectable. The proportion of erroneously uncleaved dTomato fusion constructs, that could bias sorting analysis, was negligibly low.

### Axonal Tau sorting requires the PRR2 domain but is independent of the N-terminal tail and Tau microtubule affinity

Next, in order to unravel domains or phosphorylation sites might be necessary of sufficient for axonal Tau sorting, we quantified the axonal sorting (as AEF) of all our Tau^HA^ constructs and compared it to the AEF of 0N3R-Tau^HA^.

When either the N- or C-terminal tails were missing (No-Ntail^HA^, No-Ctail^HA^), we saw sorting efficiency similar to 0N3R^HA^. Only when larger parts of the C-terminus (Nterm^HA^) or N-terminus (Cterm^HA^) were missing, Tau failed to enrich at the axonal compartment (∼5-20 % efficiency of 0N3R^HA^) (Fig. 2f). However, when the C-terminus was connected to the PRR domain, the resulting construct PRR+Cterm^HA^ showed axonal targeting like 0N3R^HA^ (Fig. 2d). The microtubule-binding domain alone without the C-terminal tail (PRR+4R^HA^) was in turn much less abundant in the axon (Fig. 2f). Significantly decreased axonal sorting was observed when Tau was lacking the repeat domains completely (no4R^HA^). The constructs only3R^HA^ and only4R^HA^ (Fig. 2f) were not enriched and not further investigated due to the lacking evidence for considerable protein levels (Fig. S3a-d).

Remarkably, the changes in dendritic enrichment (DEF) of truncated Tau^HA^ constructs followed a similar pattern to those in axonal sorting (Fig. 2g). Direct correlation of the axonal and dendritic enrichment (AEF/DEF ratio, Fig. S6b) revealed that most constructs have no changes in axon to dendrite enrichment, compared to 0N3R^HA^. Only for Cterm^HA^ and noPRR2^HA^, the AEF/DEF ratio was significantly decreased, indicating axon-specific impairments of intracellular sorting for those constructs (Fig. S6b). The nuclear levels were not changed for any truncated Tau^HA^ construct except for Nterm^HA^, likely due to the somatic inclusion formation (Fig. 2h). Previous reports claimed that stronger microtubule binding is linked to Tau missorting while weak microtubule affinity does not cause impaired axonal sorting (Iwata et al., 2019). In order to test this hypothesis with our data sets, we correlated the axonal enrichment (AEF) of all truncated Tau^HA^ constructs with their microtubule affinity (Gustke et al., 1994). In contrast to literature, the axonal enrichment did not decrease but increase with higher microtubule affinity (Fig. S6c). We observed the same positive correlation when using the AEF/DEF ratio instead of the AEF alone (Fig. S6d).

Then, we analyzed the intracellular distribution of AT8-mutant Tau^HA^ constructs. The axonal sorting was equally efficient for all constructs, regardless whether they were carrying pseudophosphorylation or dephosphorylation mutations of AT8. Only the completely pseudo-dead AT8-allA^HA^ construct was slightly higher axonally enriched than 0N3R^HA^ (Fig. 3e). In the dendrites, the same AT8-allA^HA^ construct and S199A^HA^ showed modest increase while the dendritic sorting of all other constructs was not changed, compared to 0N3R^HA^ (Fig. 3f). The nuclear enrichment was not different from 0N3R^HA^ for any of the AT8-mutant Tau^HA^ constructs (Fig. 3g).

For our KXGS-, and double-mutant Tau^HA^ constructs, the sorting seemed to become less efficient with rising number of modified residues in both cases, exchange by glutamate or alanine (Fig. 4e). The enrichment gradually declined from 1xKXGA^HA^ to 3xKXGA^HA^, and from 1XKXGE^HA^ to 2xKXGE^HA^ to 3XKXGE^HA^ (Fig. 4e), without reaching statistical significance, though. For AT8-KXGE^HA^ and AT8-KXGA^HA^, which harbor glutamate and alanine substitutions at both motifs, respectively, the levels were similar to the mutants without AT8, which is in line to the missing effect of AT8 on axonal Tau sorting (Fig. 3e). The dendritic enrichment of the KXGS-mutant constructs was decreased in a pattern similar to the axonal enrichment (Fig. 4f). Accordingly, the AEF/DEF ratio was unchanged for all constructs (Fig. S6b). for 3xKXGE^HA^. This suggests that the weak neuronal polarization is caused by a common axon-unspecific mechanism. The nuclear enrichment of all KXGS- and double-mutant Tau^HA^ constructs was not different from 0N3R^HA^ (Fig. 4g).

Of note, the intracellular sorting behavior of endogenous MAP2 was not changed upon expression of any truncated or AT8/KXGS-modified Tau^HA^ construct regarding axonal (Figs. S3g,4f,5g), dendritic (Figs. S3h,4g,5h) or nuclear sorting (Figs. S3i,4h,5i).

Taken together, we found decreased axonal targeting of the truncated protein halves Nterm^HA^ and Cterm^HA^, the repeat domain-lacking no4R^HA^, and to a lesser extent of the microtubule-binding domain PRR+3R^HA^ and PRR2-lacking noPRR2^HA^. Comparison with dendritic enrichment of all constructs revealed that only Cterm^HA^ and noPRR2^HA^ showed axon-specific missorting while both axonal and dendritic targeting was impaired for the other constructs, pointing at somatic accumulation independent of axon-specific sorting mechanisms. In contrast to literature, high microtubule affinity was not linked to impaired axonal sorting for our truncated constructs. For all our AT8-mutant Tau^HA^ constructs, axonal enrichment was equally efficient regardless of the mimicry of phosphorylation or dephosphorylation. Mutations of the KXGS residues seemed to decrease both axonal and dendritic sorting with increasing number for both glutamate and alanine but significant results are missing. Notably, the nuclear localization was not changed for any construct of our Tau^HA^ library except for Nterm^HA^, likely due to its somatic accumulation in puncta-like inclusions.

### TurboID proximity labeling is suitable for analyzing axonal binding partners of BirA-coupled Tau^HA^

Next, we aimed to unravel Tau interactions that are critical for axonal sorting. For this, we used the recently developed TurboID proximity labelling, which is based on the promiscuous biotin ligase BirA (Branon et al., 2018; Cho et al., 2020). After generating pUltra-dox vectors with HA-tagged fusion constructs of BirA and either 0N3R or the non-sorting Nterm, we produced lentiviruses and delivered them into 6-weeks-old *MAPT*-KO iPSC-neurons. After doxycycline-induced expression for 12-13 days, we applied biotin for 20 minutes to achieve efficient labelling of proteins in close proximity to BirA, and neurons were fixed or harvested immediately (Fig. 5a). The 0N4R isoform was included to check isoform-specific interaction differences.

**Fig. 5:**
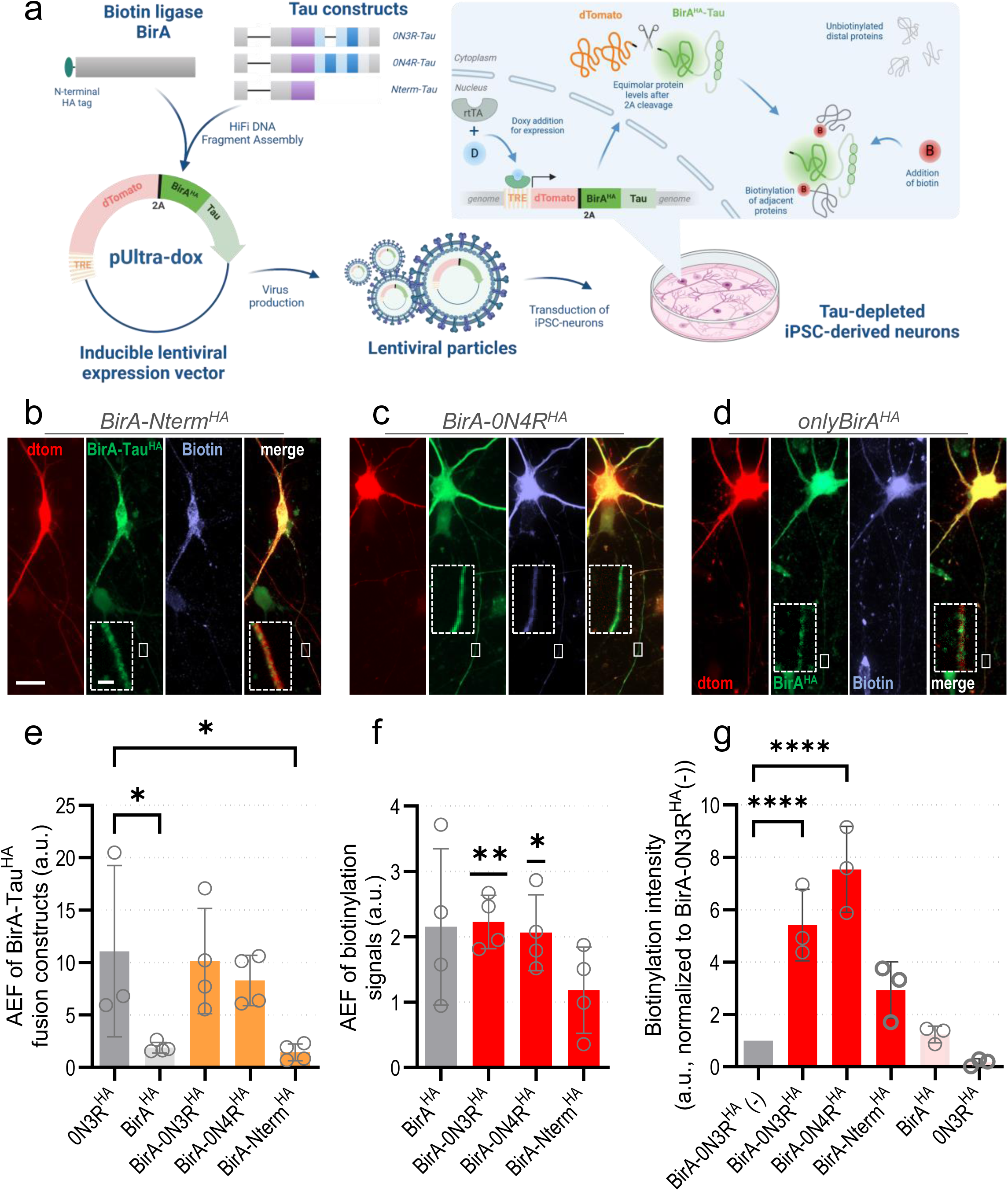
Generation & characterization of BirA-Tau fusion constructs. (a) Workflow of BirA-Tau^HA^ fusion construct generation. Coding sequences of BirA^HA^ and the Tau isoforms 0N3R and 0N4R or Nterm-Tau were cloned into the lentiviral transfer vector pUltra-dox in frame with the 2A-coupled reporter dTomato under regulation of a tetracycline response element (TRE). Virus particles were produced, harvested from HEK293T cells, and transduced to differentiated *MAPT-KO* iPSC-neurons (see Fig. S1a), resulting in undirected genomic integration of the expression cassette. Addition of doxycycline (doxy) 24 h after transduction leads to binding of the reverse tetracycline-activated transactivator (rtTA) to the TRE and equimolar expression of BirA-Tau^HA^ fusion constructs and the reporter protein dTomato. After 12-13 days of expression, biotin is added and induces biotinylation of adjacent proteins by BirA. (b-d) *MAPT*-KO iPSC-neurons after expression of BirA-Nterm^HA^ (c), BirA-0N4R^HA^ (d), and onlyBirA^HA^ (e) and dTomato for 12-13 days and subsequent immunostaining. The signals for BirA-Tau^HA^ or BirA^HA^ (green), biotin (blue) and dTomato (red) are shown in single channels and merged (green+red). Axonal sections are magnified (dashed boxes) in merged images. Scale bar: 20 µm, scale bar within dashed box: 2 µm. (e) Axonal enrichment (AEF) of BirA-Tau^HA^ fusion proteins in *MAPT*-KO iPSC-neurons, compared to the AEF of 0N3R-Tau^HA^ and BirA^HA^. Quantification was done for three to four independent experiments (black dots) with 8-18 cells per experiment. The bars indicate the arithmetic mean, error bars show SD. A mixed-effects model with Dunnett’s correction for multiple comparisons was performed to determine significance levels between the groups. Significance levels: * p < 0.05, ns: p ≥ 0.05. (f) Axonal enrichment (AEF) of biotin after expression of different BirA-Tau^HA^ fusion proteins in KO iPSC-neurons, compared to AEF of biotin in neurons expressing only BirA^HA^. Quantification was done for three to four independent experiments (black dots) with 8-18 cells per experiment. The bars indicate the arithmetic mean, error bars show SD. Groupwise one-sample t-tests were performed to determine the difference from undirected distribution (AEF = 1). Significance levels: * p < 0.05, ** p < 0.01, ns: p ≥ 0.05. (g) Biotinylation intensity after expression of BirA-Tau^HA^ fusion proteins in *MAPT*-KO iPSC-neurons, normalized to the signals in neurons expressing BirA-0N3R-Tau^HA^ without biotin treatment. Quantification was done for three to five independent experiments (black dots) with 8-18 cells per experiment. The bars indicate the arithmetic means, error bars show SD. A mixed-effects model with Dunnett’s correction for multiple comparisons was performed to determine significance levels between the groups. Non-normalized values were used to perform statistical analysis. Significance levels: * p < 0.05, ** p < 0.01, ns: p ≥ 0.05.

First, we confirmed proper expression of BirA-Tau^HA^ proteins via Western blot analysis (Fig. S10a) with only a small proportion of uncleaved side products (Fig. S7a). Importantly, the axonal enrichment of BirA-0N3R^HA^ and BirA-0N4R^HA^ was similar to that of uncoupled 0N3R^HA^ in immunostained neurons, while sorting of BirA^HA^ and BirA-Nterm^HA^ was, as expected, significantly less (Fig. 5b-d,e). For BirA-Nterm^HA^, we observed the puncta-like distribution typical for Nterm^HA^ (Fig. 5b, see also Fig. 2c)(Bell et al., 2021). The moderate but visible axonal enrichment of biotinylation signals (Fig. 5c,f) showed successful axonal diffusion of biotin during 20 minutes treatment. Longer incubation times were not feasible to prevent excessive unspecific labelling (Cho et al., 2020). Quantification of biotin signals in neurons expressing BirA-0N3R^HA^ without biotin application determined only basal biotinylation by BirA (Fig. 5g). As for our Tau library, we examined the correlation of axonal enrichment and protein levels for all BirA-Tau^HA^ constructs with all results being negative (Fig. S7c). We also did not find any differences in the relative BirA-Tau^HA^ protein levels, but a decrease for BirA^HA^ only (Fig. 7d).

All in all, we could validate that the recently developed TurboID proximity labeling is suitable to study interactions of axonally enriched proteins like Tau, as BirA-coupling did not disturb sorting efficiency and axonal biotinylation was visible.

### Key regulators of pre- and postsynaptic plasticity bind specifically to BirA-0N4R^HA^

In the next step, we isolated protein from *MAPT*-KO iPSC-neurons cultures after expression of either sample or control constructs for 12-13 days and biotin application. Biotinylated proteins were enriched with avidin-coated magnetic beads, trypsin-digested and measured with liquid chromatography-coupled mass spectrometry (LC-MS/MS) for further interactome analysis (Fig. 6a, see methods for details).

**Fig. 6:**
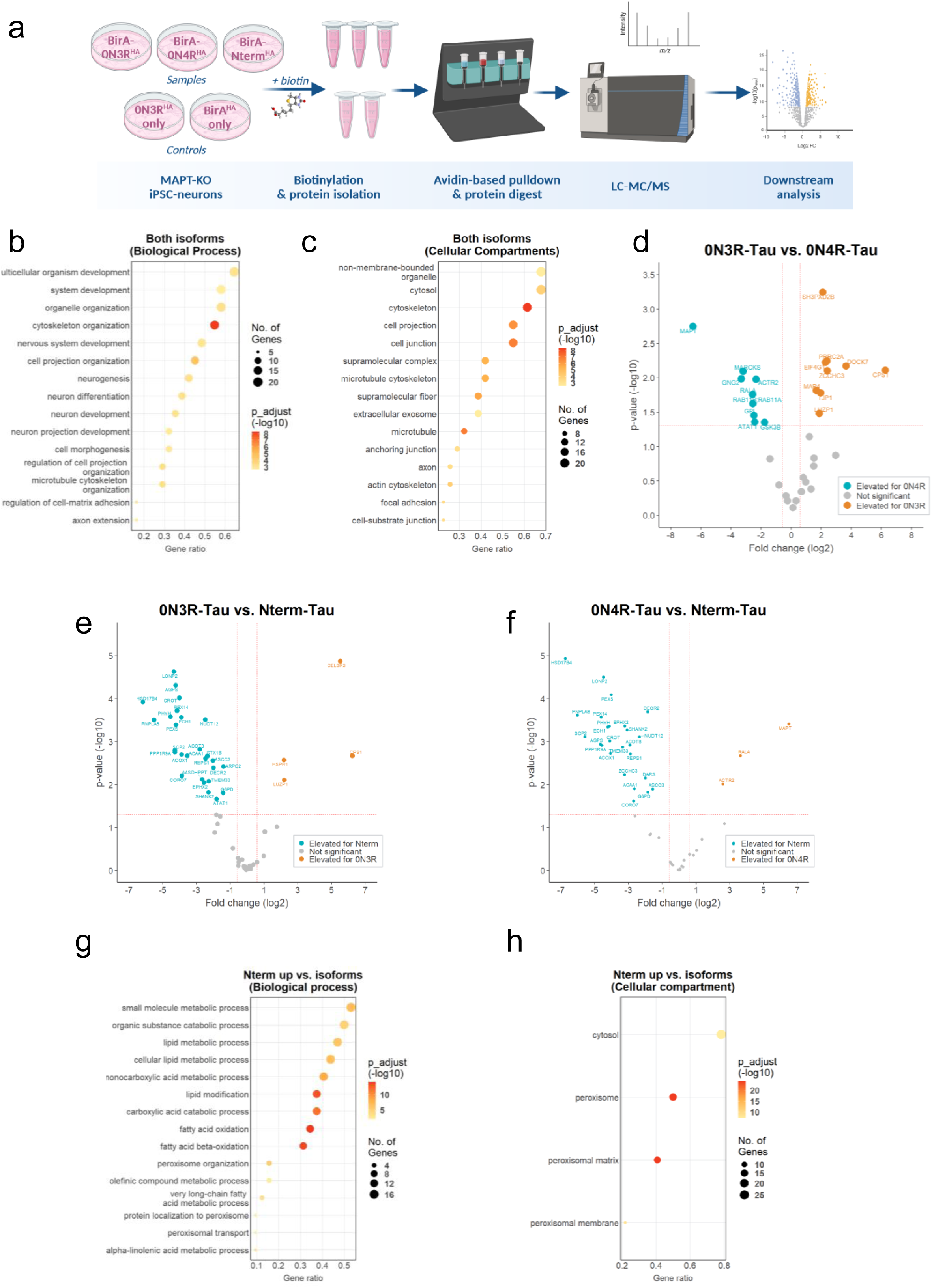
TurboID-based interactomics study of 0N3R- & 0N4R-Tau and non-sorting Nterm-Tau. (a) Workflow for interactomics sample preparation. *MAPT*-KO iPSC-derived neurons were differentiated and transduced with BirA-Tau^HA^ constructs as described (see Fig. 5a, S1a). After treatment with 500 µM biotin for 20 min, protein was isolated and incubated with avidin-coated magnetic beads for enrichment of biotinylated proteins. Enriched proteins were digested and examined with data-independent analysis via LC-MS/MS. Resulting data were used for downstream interactome analyses. (b,c) Gene ontology (GO) term analysis of interaction partners of 0N3R and 0N4R after filtering of background targets, performed with gProfiler (see methods). The GO terms for biological process (b) and cellular compartment (c) are sorted by the number of matched genes. The dot size represents the number of genes, the colour indicates the adjusted p-value. (d) Interactome comparison of the BirA-0N3R^HA^ vs. BirA-0N4R^HA^ with p-values (-log_10_) and fold-changes (log_2_) of the label-free quantification (LFQ) after filtering of background targets. Proteins enriched for 0N3R-Tau (thresholds: p-value > 1.3, fold-change < −0.58) are marked blue, proteins enriched for 0N4R-Tau (thresholds: p-value > 1.3, fold-change > 0.58) are marked orange. Gene IDs of all different interactors are given. (e,f) Interactome comparison of BirA-0N3R^HA^ vs. BirA-Nterm^HA^ (e) and BirA-0N4R^HA^ vs. BirA-Nterm^HA^ (f) with p-values (-log_10_) and fold-changes (log_2_) of the label-free quantification (LFQ), after filtering of background targets. Proteins enriched for 0N3R-Tau or 0N4R-Tau (thresholds: p-value > 1.3, fold-change < −0.58) are marked blue, proteins enriched for Nterm-Tau (thresholds: p-value > 1.3, fold-change > 0.58) are marked orange. Gene IDs of all different interactors are given. (g,h) GO term analysis of genes from (e,f) that were enriched as BirA-Nterm^HA^ binding partner, performed with gProfiler (see methods). The GO terms for and biological process (g) and cellular compartment (h) are sorted by the number of matched genes. The dot size represents the number of genes, the colour indicates the adjusted p-value.

We identified 744 proteins in total that were detected for at least one of our BirA-Tau^HA^ constructs, BirA-0N3R^HA^, BirA-0N4R^HA^ and BirA-Nterm^HA^ (Table S1a). Prior to analysis, we excluded all proteins that were not significantly enriched (see methods for details) for at least one construct compared to both control constructs, onlyBirA^HA^ and only0N3R^HA^. After this reduction step, 62 interacting proteins were remaining (Table S1b). The protein number was smaller than in a previous study using a similar approach with Tau-APEX2 constructs (Tracy et al., 2022), probably due to the exclusion of background hits found in control constructs. However, the overlap between our data set and literature was pretty large. We compared the proteins found for the isoforms BirA-0N3R^HA^ and BirA-0N4R^HA^, which were 33 out of 62, with the study using Tau-APEX2 in the identical iPSC line (Tracy et al., 2022) and found an overlap of 23 out of 33 proteins (69.7 %) (Table 1). When we checked the data from human Tau interactome meta-analysis (Kavanagh et al., 2022), 24 out of 33 proteins (72.7 %) were found (Table 1). We then conducted a gene ontology (GO) terms analysis of all 33 isoform-related hits to further validate our data sets (Tables S7,S8). We screened for sections “biological process” and “cellular compartment”. The found Tau interactors were mainly associated with microtubule function, cytoskeletal protein binding and organization, or nucleotide and ribonucleotide binding (Fig. 6b,c). This matches with the known major interaction partners of Tau (Guo et al., 2017; Kavanagh et al., 2022). This outcome confirmed the successful enrichment of Tau-interacting proteins in our experimental setup.

**Table 1:** List of all genes that found to interact either with BirA-0N3R_HA_, for BirA-0N4R_HA_, or with both isoforms. The Uniprot ID and Protein name is given for all genes. Identical gene and protein names are indicated by ‘*’. 0N3R/0N4R interactor: ‘YES’ means that the gene was found to interact in our study, ‘-‘ means that no interaction was found. Found in Kavanagh 2022: ‘YES’ means that protein was described as Tau interactor in the meta-analysis of seven human Tau interactome studies (Kavanagh et al., 2022), ‘-‘ means that it was not described in the Kavanagh et al. (2022). The number of hits refers to how many of the seven human Tau interactome papers reported Tau interaction of the protein. Found in Tracy 2022: ‘YES’ means that Tau interaction was found in Tracy et al. (2022) using biotinylation labelling in human iPSC-neurons, ‘YES*’ means that Tau interaction was not found in Tracy et al. (2022) using biotinylation labelling but FLAG-based IP of WT-Tau. ‘-‘ means that the protein was not found in Tracy et al. (2022).

| Gene | Uniprot ID | Protein | 0N3R interactor | 0N4R interactor | Found in Kavanagh 2022 | # of hits (Kavanagh 2022) | Found in Tracy 2022 |
| --- | --- | --- | --- | --- | --- | --- | --- |
| *DCX* | *O43602* | * | YES | YES | YES | 1/7 | YES |
| *MAP2* | *P11137* | * | YES | YES | YES | 3/7 | YES |
| *CAPZB* | *P47756* | * | YES | YES | YES | 2/7 | YES* |
| *MACF1* | *Q9UPN3* | * | YES | YES | - | - | - |
| *CASK* | *O14936* | * | YES | YES | YES | 1/7 | YES* |
| *LUZP1* | *Q86V48* | * | YES | - | - | - | - |
| *MAP4* | *P27816* | * | YES | - | YES | 4/7 | YES |
| *CPS1* | *P31327* | CPSM | YES | - | YES | 1/7 | - |
| *PRRC2A* | *P48634* | PRC2A | YES | - | - | - | - |
| *EIF4G1* | *Q04637* | IF4G1 | YES | - | YES | 3/7 | YES* |
| *TJP1* | *Q07157* | ZO1 | YES | - | YES | 1/7 | YES* |
| *SH3PXD2B* | *A1X283* | SPD2B | YES | - | - | - | - |
| *MINK1* | *Q8N4C8* | * | YES | - | - | - | - |
| *DOCK7* | *Q96N67* | * | YES | - | - | - | - |
| *HSPH1* | *Q92598* | HSP110 | YES | - | YES | 3/7 | YES* |
| *FERMT2* | *Q96AC1* | FERM2 | YES | - | - | - | - |
| *HN1* | *Q9UK76* | JUPI1 | YES | - | YES | 1/7 | YES* |
| *EML4* | *Q9HC35* | EMAL4 | YES | - | YES | 1/7 | YES* |
| *ZCCHC3* | *Q9NUD5* | ZCHC3 | YES | - | - | - | - |
| *CELSR3* | *Q9NYQ7* | CELR3 | YES | - | - | - | - |
| *GPI* | *P06744* | G6PI | - | YES | YES | 3/7 | YES |
| *MAPT* | *P10636* | Tau | - | YES | YES | 5/7 | YES |
| *RALA* | *P11233* | * | - | YES | YES | 1/7 | YES* |
| *ACTR2* | *P61160* | ARP2 | - | YES | YES | 1/7 | YES* |
| *GSK3B* | *P49841* | GSK3B | - | YES | YES | 2/7 | YES* |
| *MARCKS* | *P29966* | MARCKS | - | YES | YES | 2/7 | YES* |
| *ZC2HC1A* | *Q96GY0* | ZC21A | - | YES | YES | 1/7 | YES* |
| *GNG2* | *P59768* | GBG2 | - | YES | YES | 1/7 | YES* |
| *RAB11A* | *P62491* | RB11A | - | YES | YES | 1/7 | YES* |
| *RAB11B* | *Q15907* | RB11A | - | YES | YES | 1/7 | YES* |
| *ATAT1* | *Q5SQI0* | * | - | YES | YES | 1/7 | YES* |
| *KIF21A* | *Q7Z4S6* | KI21A | - | YES | YES | 2/7 | YES |

We analyzed which of the 33 proteins interacting with either 0N3R or 0N4R were significantly enriched for one of the isoforms (Figs. 6d, S11g, Tables S2,S9). The 9 proteins that specifically interacted with 0N3R, exhibit diverse functions. Three were either directly, LUZP1 and TJP1, or indirectly, SH3PXD2B, linked to the f-actin cytoskeleton. MAP4 is a microtubule-binding protein that is structurally related to Tau as both are members of the same MAP family (Nishida et al., 2023; Tokuraku et al., 2010).

Out of the 10 specific interactors of 0N4R, at least 5 proteins are connected to pre- or postsynaptic plasticity, vesicle release and neurite growth, namely the GTPases RALA, RAB11A, RAB11B, the signaling protein MARCKS, and the synaptic serine/threonine kinase GSK-3beta (Arendt et al., 2016; Brudvig et al., 2018; Lalli, 2009; Lalli & Hall, 2005; Sultana & Novotny, 2022; Xu et al., 2014; Zempel & Mandelkow, 2014). MARCKS and GSK-3beta are both linked to CDC42 (Brudvig et al., 2018; Zhu et al., 2023), a key regulator of postsynaptic f-actin plasticity. Abnormally high CDC42 activity is observed in AD patients, and can evoke AD-like phenotypes in mice, presumably by overactivation of GSK-3beta (Zhu et al., 2023). MARCKS also plays a role at the presynapse where it controls vesicle docking to the active zone (Xu et al., 2014). The 0N4R interactors RALA and RAB11A/B also affect presynaptic function, as they interact with the exocyst complex, a membrane complex that tethers secretory vesicles to the membrane and thereby plays a fundamental role in polarized exocytosis in eukaryotic cells (Lalli, 2009; Lalli & Hall, 2005; Sultana & Novotny, 2022). The remaining 0N4R-specific interactors were either linked to cytoskeletal functions, such as the microtubule acetyltransferase ATAT1 and the f-actin-organizing protein ACTR3 or to metabolic function (GPI).

### The non-sorting BirA-Nterm^HA^ shows peroxisomal association while CELSR3 and HPSH1 are interactors specific for sorting BirA-Tau^HA^

Next, we compared the interactome of the isoforms, BirA-0N3R^HA^ and BirA-0N4R^HA^, with that of BirA-Nterm^HA^, which does not show axonal enrichment (Fig. 6e,f, Tables S10,11). The analysis revealed that the BirA-Nterm^HA^ interactome comprised several proteins that were not interacting with any of the isoforms. In contrast, only few proteins were specific to the isoforms (Table S3). We performed GO term analysis of the Nterm-specific proteins for “biological process” and “cellular compartment” (Tables S4-S6). The overwhelming number of terms, that we found, were linked to the “cellular compartments” peroxisomal membrane and peroxisomal matrix (Fig. 6h), and lipid catabolism, fatty acid oxidation, and related peroxisomal processes as main “biological processes” (Fig. 6g). These findings suggest that the puncta-like inclusions of Nterm^HA^, which we observe for both the single construct (Fig. 2c) and the BirA-Nterm^HA^ fusion construct (Fig. 5b), represent peroxisomal localization. How Nterm-Tau gets imported into the peroxisomal lumen, remains unclear. Nterm-Tau does not contain the typical peroxisomal targeting signals PTS1 and PTS2 (Brocard & Hartig, 2006; Lametschwandtner et al., 1998; Neuberger et al., 2003a, 2003b; Purdue & Lazarow, 2001), and two in silico tools (PTS1 predictor: https://mendel.imp.ac.at/pts1/, PSORTII: https://psort.hgc.jp/form2.html) predicted it as non-targeting based on the amino acid sequence. However, the specific interaction with two proteins of the PTS1-associated import complex (PEX5, PEX14) clearly hint at PTS1-mediated translocation.

Only few proteins were enriched for the sorting Tau^HA^ isoforms compared to the non-sorting Nterm-Tau^HA^ (Table S3). Out of those proteins, only the G-protein coupled receptor CELSR3 and the heat shock protein HSPH1 were enriched in both isoforms. CELSR3 is involved in axon guidance of developing corticothalamic projections, but interaction with Tau was not previously reported. HSPH1 is known to interact with Tau and the major axonal Tau phosphatase PP2A (Eroglu et al., 2010; Kavanagh et al., 2022).

In brief, these results hint at the import and accumulation of BirA-Nterm^HA^ inside the peroxisomal lumen. The few sorting-specific interaction partners are involved in axon guidance and modulation of the axonal Tau-targeting phosphatase PP2A.

## Discussion

Somatodendritic missorting is an early sign of Tau pathology in AD and related diseases (Braak & Del Tredici, 2011; Braak et al., 2011). To date, it remains largely elusive how axonal Tau sorting is mediated under healthy conditions and what leads to disruption of these processes in disease conditions. In this study, we aimed to identify domains or motifs of Tau and cellular interaction partners that are required for efficient axonal Tau sorting. Further, we wanted to unravel isoform-specific Tau interaction partners that might support the idea of different roles of human Tau isoforms in normal function and in disease development (Buchholz & Zempel, 2024).

### Correct expression of truncated and phosphorylation-mutant Tau constructs in human MAPT-KO iPSC-derived neurons

With our experimental approach, we overcame several limitations of previous studies: i) We could study sorting behavior without interference with endogenous Tau by using the recently engineered *MAPT*-knock out (KO) iPSC line (Buchholz et al., 2022), carrying a *Ngn2* transgene for enabling rapid neuronal differentiation (Miyaoka et al., 2014; Zhang et al., 2013), ii) we achieved axonal sorting of exogenous 0N3R-Tau similar to wildtype Tau by using lentiviral delivery with inducible expression (Iwata et al., 2019; Xia et al., 2016), and iii) the use of human iPSC-derived neurons instead of rodent primary neurons (Gauthier-Kemper et al., 2018; Iwata et al., 2019; Li et al., 2011; Zempel et al., 2017) diminished the problem of inter-species differences in Tau physiology, such as isoform expression pattern and interaction profiles (Kavanagh et al., 2022; Kosik et al., 1989).

Prior to analysis, we confirmed correct Tau^HA^ construct expression and relative protein levels by Western blot and immunofluorescent analysis. For all constructs used in this study, we saw negligibly low levels of Tau^HA^-dTomato fusion proteins, indicating successful cleavage of the 2A peptide (Szymczak-Workman et al., 2012). The truncated only3R^HA^ and only4R^HA^ were not detected in both assays, presumably due to proteasomal degradation or recognition by another protein quality control pathway (Amm et al., 2014; Pohl & Dikic, 2019; Ruggiano et al., 2014). Notably, the KXGS-mutant Tau^HA^ constructs exhibited a slightly smaller but distinct additional protein species that became more abundant with more mutated residues and independent of exchange to glutamate or alanine. We hypothesize that disturbed interaction with KXGS-targeting enzymes such as MARK or PP2A (Chudobová & Zempel, 2023; Gong et al., 2000) could affect the posttranslational state of adjacent residues, or that the Tau^HA^ mutants become target of proteolytic cleavage, which is known to produce similarly sized fragments under pathological conditions (Corsetti et al., 2008; Corsetti et al., 2015; Park & Ferreira, 2005). Nonetheless, we found that our Tau^HA^ constructs were largely expressed correctly after lentiviral transduction.

### Axonal Tau sorting is independent of the N-terminal tail, the C-terminal repeat domains, and the general microtubule affinity

When we tested the axonal sorting of our Tau^HA^ constructs in human *MAPT*-KO iPSC-derived neurons, we observed poor axonal sorting of both individual protein halves. Hence, either both N-terminal and C-terminal parts of Tau are required for axonal sorting, or the main axonal traffic checkpoint, the axon initial segment (AIS) (Leterrier, 2016, 2018; Rasband, 2010), prevents undisturbed axonal transit. For the N-terminal half, we confirmed findings from human SH-SY5Y-derived neurons and mouse primary neurons (Bell et al., 2021). In all three models, the N-terminal half accumulated in distinct puncta-like inclusions (Bell et al., 2021), which were absent for all other studied constructs. Of note, some of the investigated constructs (e.g. PRR+3R-Tau or no4R-Tau) showed similar changes in axonal and dendritic sorting, hinting at confounding processes that cannot be explained with axonal sorting mechanisms.

Previous studies with GFP-coupled Tau claimed that the N-terminal tail of Tau to be critical for axonal sorting. A Tau fragment lacking the N-terminal tail failed to interact with the axonal membrane proteins annexin A2 and A6 and was abnormally redistributed to the soma (Gauthier-Kemper et al., 2018). We could not confirm the role of the N-terminal tail as our noNTail-Tau construct showed efficient axonal sorting. This result might not counterargue the previous findings, as we determined the net axonal enrichment instead of only retrograde protein flux. Thus, the effects seen previously (Gauthier-Kemper et al., 2018) could be compensated by enhanced anterograde transport. Indeed, the same effect was observed for 2N4R-Tau with complete pseudophosphorylation of KXGE: The 2N4R-Tau mutant was axonally enriched similar to wildtype 2N4R-Tau although it showed increased retrograde diffusion (Li et al., 2011; Zempel et al., 2017). Another recent study revealed that Tau shows weak axonal sorting when the complete PRR2 domain is dephosphorylation-mimetic (Iwata et al., 2019). Since the microtubule affinity of this mutant Tau was modestly increased, and a 4R-depleted Tau mutant with less microtubule binding was properly sorted, the authors claim that tight microtubule binding impairs axonal Tau sorting.

In our study, the phosphorylation-mimetic AT8- and KXGS mutants (both Ser to Glu) showed no deficits of axon-specific sorting, supporting the idea of efficient sorting of highly diffusible Tau (Iwata et al., 2019; Zempel et al., 2017). However, the role of Tau microtubule affinity in axonal sorting appears questionable due to several observations: The phosphorylation-dead AT8 mutants (Ser to Ala) and KXGA mutants were sorted equally well in our *MAPT*-KO iPSC-derived neurons, contradicting the previous outcomes (Iwata et al., 2019). Further, the altered microtubule affinity of the phosphorylation-dead PRR2 mutant, suggested to be responsible for its somatic retention, changes the proportion of bound microtubules only marginally compared to wildtype Tau (Iwata et al., 2019), possibly due to low levels of basal PRR2 phosphorylation. When we depleted the entire PRR2 domain, thus also decreasing its microtubule affinity (Gustke et al., 1994; Iwata et al., 2019), the resulting construct showed poor axonal enrichment. An effect that was in fact also found in the former study (Iwata et al., 2019). Depletion of the C-terminal repeat domains, in turn, had no effect on axon-specific Tau sorting in our and previous studies (Iwata et al., 2019). To summarize, highly diffusible Tau can be sorted efficiently (KXGE mutants, 4R-lacking construct)(Iwata et al., 2019; Zempel et al., 2017) or inefficiently (PRR2-lacking, our study, (Iwata et al., 2019), and also microtubule-affine Tau can be sorted efficiently (KXGA mutants, AT8 Ser->Glu mutants, our study) or inefficiently (PRR2 Ser->Glu construct (Iwata et al., 2019). Thus, we postulate that both the C-terminal repeat domains and the microtubule affinity of Tau play no key role in the axonal Tau sorting process.

### Axonal Tau sorting depends on the PRR2 domain independent of AT8 phosphorylation

In contrast, certain residues or motifs of the PRR2 domain appear critical for the axonal sorting of Tau. The unaltered sorting of all AT8 mutants makes this motif unlikely to be involved in this mechanism. However, sorting analysis of an AT8-depleted Tau construct might reveal whether AT8 residues are involved independent of the phosphorylation state. It would be also interesting to whether a construct with all PRR2 being pseudo-phosphorylated resembles the results of the used phosphorylation-dead construct (Iwata et al., 2019). If yes, this would further confirm the irrelevance of microtubule affinity and suggest a mechanism that depends on PRR2-specific interaction partners or the exact wildtype amino acid sequence. The PRR was shown to contribute to the microtubule-binding geometry of Tau by binding the outer tubulin surface (Amos, 2004; Kar et al., 2003).

These results also question the idea that hyperphosphorylation of the AT8 and KXGS motifs, which are hallmarks of pathological Tau missorting (Zempel & Mandelkow, 2014), is causative for the abnormal cellular distribution. Our results rather hint at hyperphosphorylation of AT8 and KXGS residues upon somatodendritic missorting or a combinatorial effect of AT8, KXGS, and other phosphorylation sites that we did not address.

### 0N4R-Tau interacts with regulators of synaptic plasticity that are involved in AD pathogenesis

The human Tau isoforms differ in their intracellular localization (Bachmann, Bell, et al., 2021; Buchholz et al., 2022; Zempel et al., 2017), and co-immunoprecipitation (Co-IP) experiments in rodent have revealed distinct binding partners suggesting isoform-specific roles in cellular Tau function (Liu et al., 2016). Comparable data are lacking for human Tau, although there is evidence for differential roles of human Tau isoforms in both health and disease (Buchholz & Zempel, 2024). Thus, we next looked for interaction partners that are specific for individual Tau isoforms, with the aim to find evidence for distinct functional roles in health and disease.

We employed the proximity labelling technique TurboID, which is based on promiscuous biotinylation by the biotin ligase BirA (Branon et al., 2018; Cho et al., 2020), to find specific interactors of the 0N3R-Tau and 0N4R-Tau in our *MAPT*-KO iPSC-derived neurons. Importantly, we confirmed that coupling with BirA did not compromise the sorting efficiency of both isoforms as shown here. In previous studies, interaction partners were consistent between uncoupled Tau and Tau bound to the peroxidase APEX2 (Kavanagh et al., 2022; Tracy et al., 2022), indicating that our approach was suitable to identify physiologically relevant interactors. Around 60 % of interactors that we identified for both Tau isoforms were also found in a recent study using biotinylation labelling in the same neuronal model (Tracy et al., 2022), and more than 75 % were described in the Tau interactome meta-analysis (Kavanagh et al., 2022). Consistently, our gene ontology (GO) term analysis confirmed that the binding partners were mainly associated with cytoskeletal organization, nucleotide and ribonucleotide binding.

For both isoforms, we found specific interaction with microtubule-associated proteins. 0N3R-Tau interacted with MAP4, which is structurally similar to Tau and controls microtubule branching in neurons (Nishida et al., 2023; Tokuraku et al., 2010). 0N4R-Tau showed association with ATAT1, the enzyme responsible for acetylation of stable microtubules (Iuzzolino et al., 2024; Li & Yang, 2015). This could suggest that 0N4R-Tau promotes microtubules stability while 0N3R-Tau rather enhances plasticity, in line with the previous claim of labile plasticity being the key benefit of Tau-microtubule binding (Qiang et al., 2018) Overexpression of individual isoforms in SH-SY5Y cells did not result in changed microtubule stability or growth (Bachmann, Bell, et al., 2021). Previous studies showed that Tau binds f-actin with the repeat domains (Correas et al., 1990; Elie et al., 2015). Accordingly, we saw different f-actin-related binding partners for 0N3R-Tau (LUZP1, TJP1) and 0N4R-Tau (ACTR2, MARCKS).

The 0N4R-specific signaling protein MARCKS regulates various cellular functions (Fanning et al., 1998; Furuse et al., 1994), and plays a crucial role in regulating synaptic plasticity (Brudvig et al., 2018). At the postsynapse, MARCKS controls dendrite growth and spine formation via activation of the CDC42 pathway (Brudvig et al., 2018). CDC42 is a membrane-bound Rho GTPase that was found to be upregulated in AD patients (Ying et al., 2022; Zhu et al., 2023). Pathological CDC42 overactivation can induce f-actin breakdown, synapse defects, and Tau hyperphosphorylation via recruitment of the Tau kinase GSK-3β (Zhu et al., 2023). Strikingly, we identified also GSK-3β as 0N4R-specific interactor in our analysis. Since GSK-3β targets the PRR2 and C-terminal tail domains of Tau (Zempel & Mandelkow, 2014), this isoform-specific interaction cannot be explained with the additional repeat domain. These findings rather suggest that 0N4R-Tau (or 4R isoforms in general) is both an upstream regulator of CDC42-mediated postsynaptic function and a downstream effector of CDC42-induced AD-related spine dysfunction.

Further, we found 0N4R-specific interactors that control presynaptic function. The multifunctional signaling protein MARCKS controls not only postsynaptic processes, but affects also secretion of RAB10-positive vesicles that are involved in axon development (Xu et al., 2014). Another binding partner, the Ras-related GTPase RALA controls vesicle trafficking by interacting with the exocyst (Lalli, 2009; Lalli & Hall, 2005), an octameric membrane complex that plays a fundamental role in polarized exocytosis in all eukaryotic cells by tethering secretory vesicles to the membrane (Hsu et al., 1998). In neurons, exocyst-dependent exocytosis regulates neurite outgrowth, synaptic receptor transport and neuronal polarity (Lalli, 2009; Lalli & Hall, 2005; Mehta et al., 2005; Sans et al., 2003; Vega & Hsu, 2001). Two more 0N4R-interactors, RAB11A and RAB11B, are closely connected to the exocyst (Sultana & Novotny, 2022), as they interact with multiple complex proteins and mediate vesicle tethering, as shown in yeast (Das & Guo, 2011). Of note, RAB11 has been connected to AD pathogenesis as it controls the endosomal recycling of beta-secretase (Li et al., 2012). Reduction of RAB11 levels promoted BACE-mediated APP cleavage and accumulation of pathogenic amyloid-β (Sultana & Novotny, 2022; Udayar et al., 2013).

Previous studies showed that Tau binds to presynaptic vesicles and proteins of the active zone and contributes to synapse dysfunction observed in AD (McInnes et al., 2018; Tracy et al., 2022; Zhou et al., 2017). Here, we provide the first evidence that presynaptic interactions are isoform-specific and that 0N4R-Tau specifically binds signaling proteins modulating the exocyst-dependent exocytosis and RAB11 protein function.

### The Tau N-terminal half accumulates in peroxisomes while axonal Tau specifically binds the PP2A activator HSP110

Next, we compared the interactome of the Tau isoforms 0N3R and 0N4R with that of a construct which shows no axonal sorting. The identification of axonal sorting-specific Tau interactors is crucial to understand the Tau sorting process and to develop targeted treatments that could restore the axonal targeting under disease-related missorting conditions.

For the somatically retained N-terminal fragment, we found numerous specific binding partners and the majority of them were peroxisomal proteins. Association of Tau with the peroxisome, the major organelle for metabolizing fatty acids and other lipids (Eckert & Erdmann, 2003; Wanders & Waterham, 2006), were not described for both healthy and disease conditions (Kavanagh et al., 2022). When Tau undergoes proteolytic cleavage under disease conditions, fragments similar to our constructs are generated (Guo et al., 2017). The NH2 fragment (aa26-230) was enriched in synaptic mitochondria (Corsetti et al., 2008; Corsetti et al., 2015), while a similar fragment (aa45-230) showed random cellular distribution (Ferreira & Bigio, 2011; Park & Ferreira, 2005). We found that Nterm-Tau binds to the membrane receptor peroxin 5 (PEX5) and another component of the PEX5-associated complex (Platta & Erdmann, 2007). This complex mediates the import of most peroxisomal proteins (Platta & Erdmann, 2007). How Nterm-Tau interacts with the PEX5 complex remains unclear since the construct lacks features of the C-terminal recognition sequence (Brocard & Hartig, 2006; Lametschwandtner et al., 1998; Neuberger et al., 2003a, 2003b) and it was classified as non-targeting for peroxisomes by two in silico prediction tools (see results).

We found two proteins that were enriched for both isoforms compared to the non-sorting Nterm-Tau, CELSR3 and HSPH1. All other found proteins were either enriched only for 0N3R-Tau or 0N4R-Tau, indicating axonal sorting-independent effect as both isoforms show roughly equal sorting efficiency (Bachmann, Bell, et al., 2021). The G-protein coupled receptor CELR3, encoded by the CELSR3 gene, is involved in axon guidance via the WNT/PCP pathway (Qu et al., 2014; Zhou et al., 2008). Loss of synaptic CELR3 impairs corticothalamic projections and reduces glutamatergic synapse formation (Feng et al., 2016; Thakar et al., 2017). Interaction with Tau was not described previously (Kavanagh et al., 2022). In contrast, the heat shock protein HSP110, encoded by *HSPH1*, is a well-described interactor of axonal Tau (Kavanagh et al., 2022). HSP110 regulates the activity of PPA2 (Eroglu et al., 2010), the major axonal Tau phosphatase (Arendt et al., 2016), and HSP110 depletion causes Tau hyperphosphorylation of axonal Tau (Eroglu et al., 2010). Thus, HSP110 could be a key factor for efficient axonal retention of Tau by promoting microtubule binding, independent of the isoform. Our findings from axonal sorting of highly diffusible Tau constructs, however, question the contribution of retrograde retention to the net axonal enrichment.

Taken together, the investigation of sorting-specific interactors provides evidence that the non-sorting Nterm-Tau gets imported to the peroxisomal lumen where it accumulates. It remains elusive how Nterm-Tau gets recognized by the import machinery and why Nterm-Tau is sequestered inside an organelle that is not involved in cellular protein quality control (Eckert & Erdmann, 2003; Wanders & Waterham, 2006). The number of binding partners that we identified as specific for sorting Tau constructs was low. Out of the two proteins, the heat chaperone HSP110 affects axonal PP2A activity and could directly regulate the axonal retention of Tau.

## Conclusion

Somatodendritic missorting of Tau is an early and crucial event in AD-related Tau pathology. However, the knowledge about Tau-intrinsic factors and Tau binding partners that mediate the axonal Tau sorting is still sparse. Here, we provide evidence that the N-terminal tail of Tau and the general microtubule affinity of the protein play a minor role in axonal Tau enrichment in human mature iPSC-derived neurons that do not express endogenous Tau. In turn, we postulate that the PRR2 domain is a major regulator of axonal Tau sorting by a mechanism independent of AT8 phosphorylation. While the non-sorting Nterm-Tau accumulates inside peroxisomes, we identified the PP2A activator HSP110 as an axonally sorted Tau binding partner, implying a potential role of PP2A activity in axonal Tau retention.

We provide evidence for isoform-specific functions in health and disease. We analyzed the differences of the isoform-specific interactome between 0N3R- and 0N4R-Tau, and found interaction with pathways that are involved in pre- and postsynaptic plasticity and associated with AD-related synaptic dysfunction, such as the CDC42 pathway or RAB11 proteins. This underlines the differential role of Tau isoforms both in basic physiological function and Tau-associated disease development, but more research is needed to understand these roles in more detail.

## Supporting information

Supplemental script 1

## Acknowledgments

Animals were obtained from the CMMC animal facility and the CECAD in vivo research facility (both Cologne, Germany). All imaging and axotomy experiments were performed at and supported by the CECAD imaging facility. Cell culture work with human iPSCs and iPSC-derived neurons was performed in the CMMC iPSC facility. This work was funded by the Else-Kröner-Fresenius Stiftung, Deutsche Forschungsgemeinschaft (DFG), and supported by a doctoral fellowship of the Studienstiftung des deutschen Volkes. The authors declare that they have no competing interests. We thank Daniel Adam for supporting the bioinformatical data analysis.

## Author contributions

Study design: MBS, HZ. Experimental work: MBS, JK. Methodological support: SB. Data analysis and interpretation: MBS, HZ. Manuscript writing: MBS, HZ. Manuscript proofreading: SB, JK.

