## Supplemental Figures for "Axonal Tau sorting depends on the PRR2 domain and 0N4R-specific interactions hint at distinct roles of Tau isoforms in synaptic plasticity"

Suppl.Fig. 1

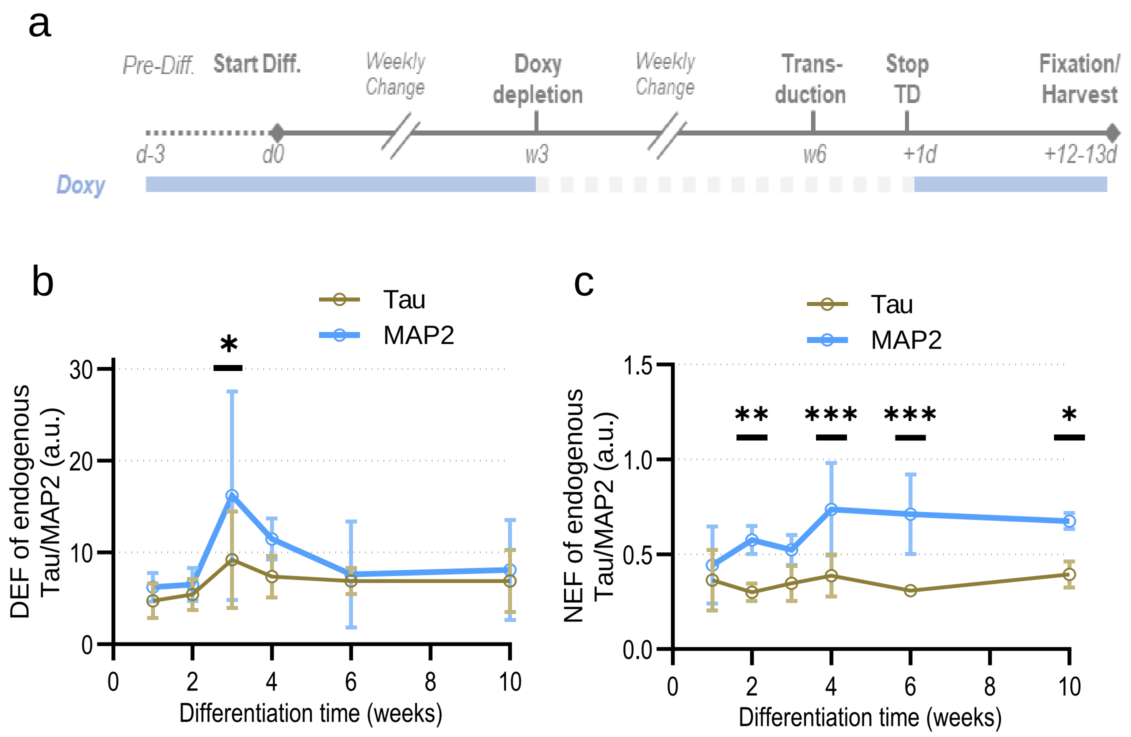
a
Pre-Diff.
Start Diff.
Weekly Change
Doxy depletion
Weekly Change
Trans-duction
Stop TD
Fixation/ Harvest
d-3
d0
w3
w6
+1d
+12-13d
Doxy
b
Tau
MAP2
DEF of endogenous Tau/MAP2 (a.u.)
Differentiation time (weeks)
c
Tau
MAP2
NEF of endogenous Tau/MAP2 (a.u.)
Differentiation time (weeks)

Suppl.Fig. 2

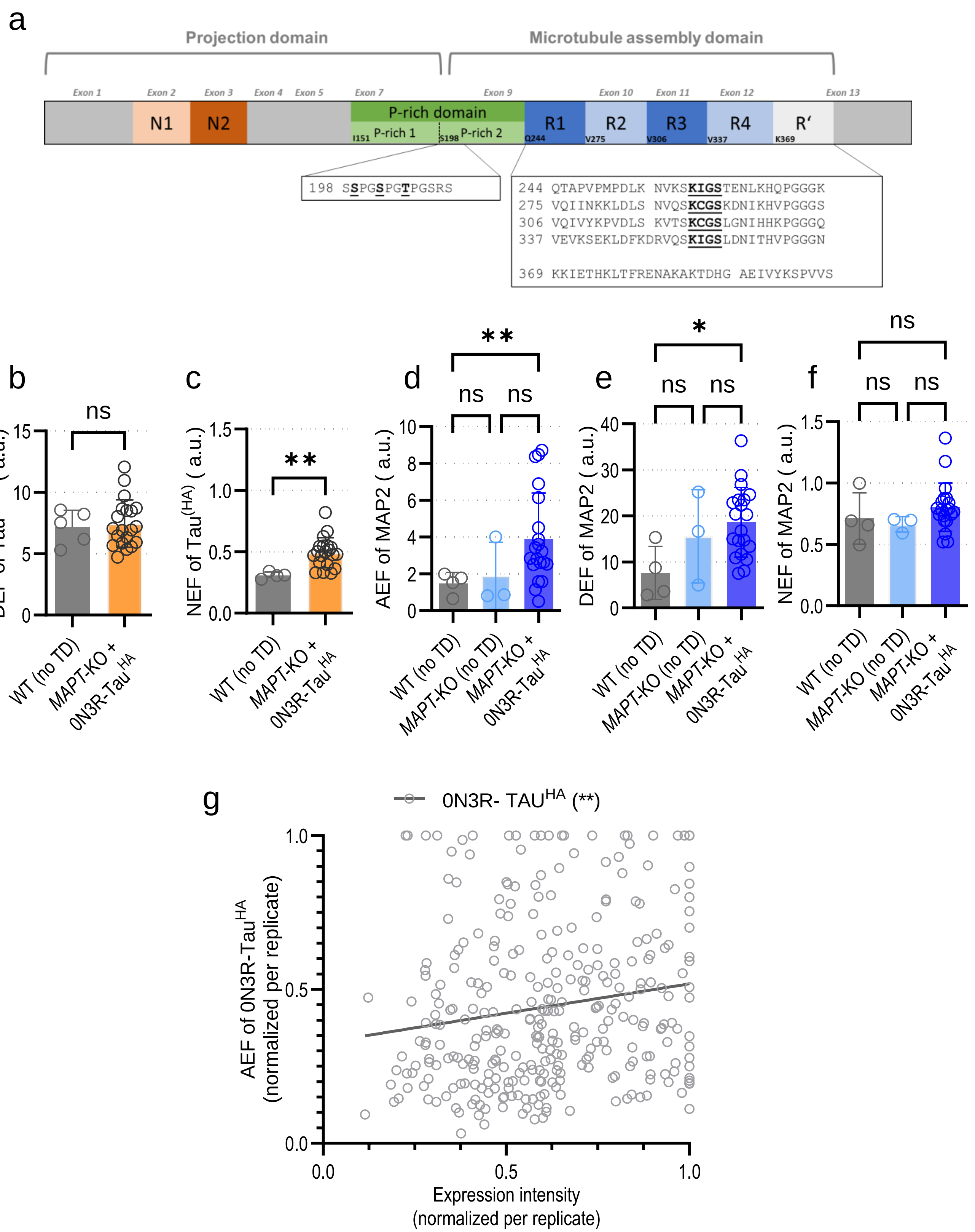
a
Projection domain
Microtubule assembly domain
Exon 1
Exon 2
Exon 3
Exon 4
Exon 5
Exon 7
Exon 9
Exon 10
Exon 11
Exon 12
Exon 13
N1
N2
P-rich domain
P-rich 1
P-rich 2
I151
S198
R1
R2
R3
R4
R'
Q244
V275
V306
V337
K369
198 SSPGSPGTPGSRS
244 QTAPVPMPDLK NVKSKIGSTENLKHQPGGGK
275 VQIINKKLDLS NVQSKCGSKDNIKHVPGGGS
306 VQIVYKPVDLS KVTSKCGSLGNIHHKPGGGQ
337 VEVKSEKLDFKDRVQSKIGSLDNITHVPGGGN
369 KKIETHKLTFRENAKAKTDHG AEIVYKSPVVS
b
DEF of Tau(HA) ( a.u.)
ns
WT (no TD)
MAPT-KO + 0N3R-Tau HA
c
NEF of Tau(HA) ( a.u.)
**
d
AEF of MAP2 ( a.u.)
**
ns
ns
MAPT-KO (no TD)
e
DEF of MAP2 ( a.u.)
*
f
NEF of MAP2 ( a.u.)
g
0N3R- TAU HA (**)
AEF of 0N3R-Tau HA (normalized per replicate)
Expression intensity (normalized per replicate)

Suppl.Fig. 3

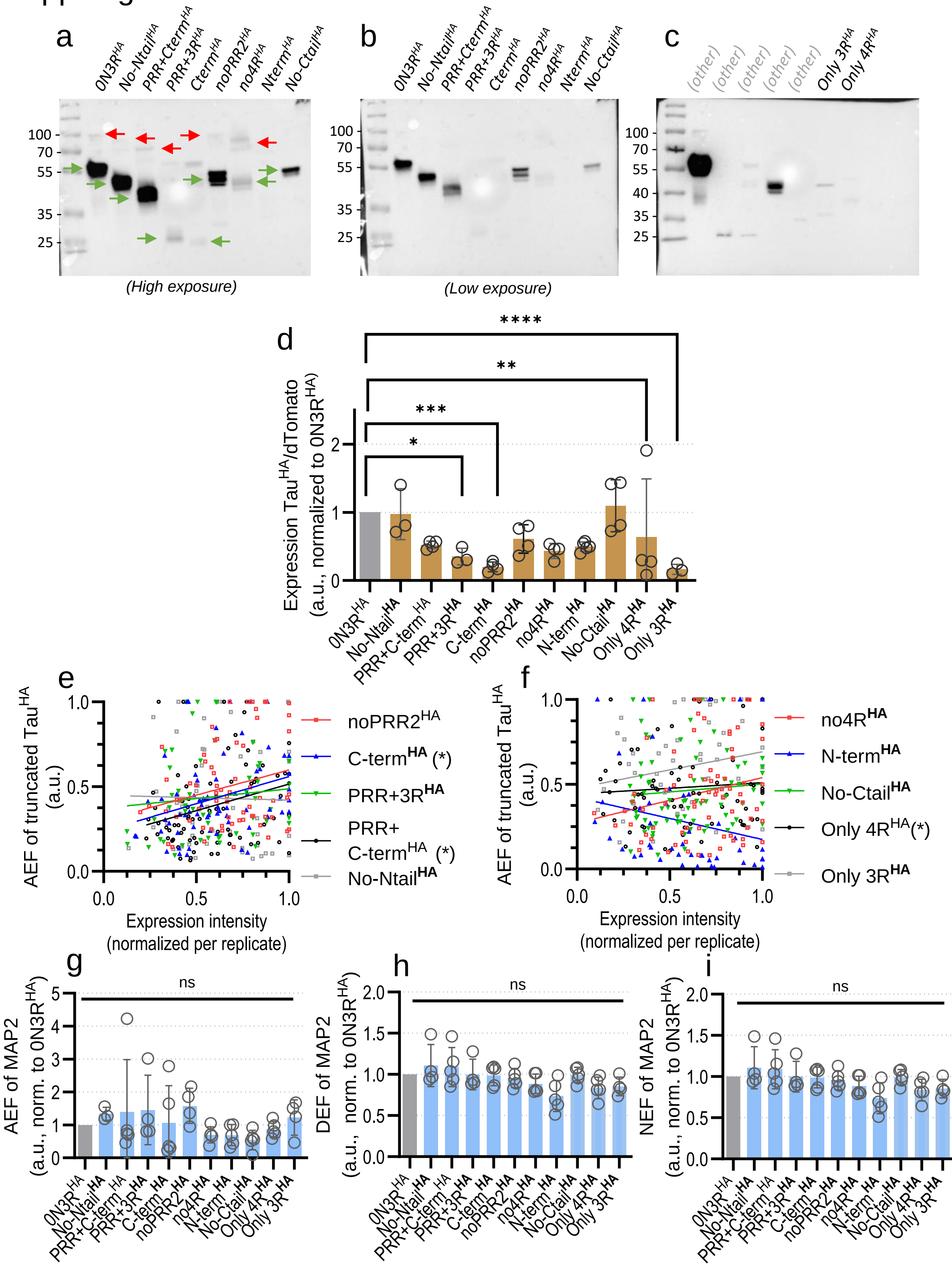
a
0N3R^HA
No-Ntail^HA
PRR+Cterm^HA
PRR+3R^HA
Cterm^HA
noPRR2^HA
no4R^HA
Nterm^HA
No-Ctail^HA
100
70
55
35
25
(High exposure)
b
(Low exposure)
c
(other)
Only 3R^HA
Only 4R^HA
40
d
Expression Tau^HA/dTomato (a.u., normalized to 0N3R^HA)
****
**
***
*
0N3R^HA
No-Ntail^HA
PRR+C-term^HA
PRR+3R^HA
C-term^HA
noPRR2^HA
no4R^HA
N-term^HA
No-Ctail^HA
Only 4R^HA
Only 3R^HA
e
AEF of truncated Tau^HA (a.u.)
noPRR2^HA
C-term^HA (*)
PRR+3R^HA
PRR+ C-term^HA (*)
No-Ntail^HA
Expression intensity (normalized per replicate)
f
no4R^HA
N-term^HA
No-Ctail^HA
Only 4R^HA(*)
Only 3R^HA
g
AEF of MAP2 (a.u., norm. to 0N3R^HA)
ns
h
DEF of MAP2 (a.u., norm. to 0N3R^HA)
i
NEF of MAP2 (a.u., norm. to 0N3R^HA)

Suppl.Fig. 4

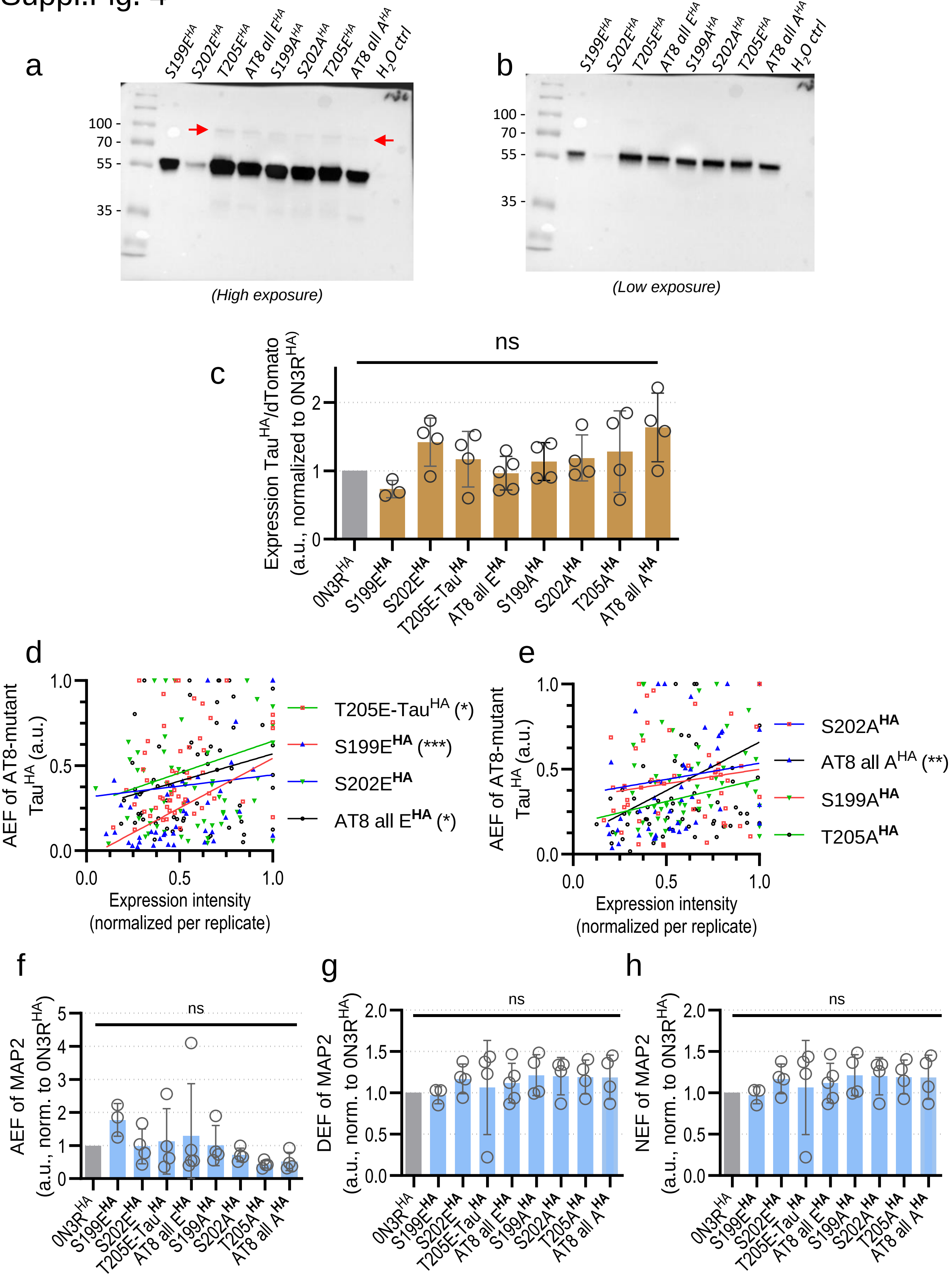

a
S199E^HA
S202E^HA
T205E^HA
AT8 all E^HA
S199A^HA
S202A^HA
T205E^HA
AT8 all A^HA
H2O ctrl
100
70
55
35
(High exposure)
b
S199E^HA
S202E^HA
T205E^HA
AT8 all E^HA
S199A^HA
S202A^HA
T205E^HA
AT8 all A^HA
H2O ctrl
100
70
55
35
(Low exposure)
c
ns
Expression Tau^HA/dTomato (a.u., normalized to 0N3R^HA)
0N3R^HA
S199E^HA
S202E^HA
T205E-Tau^HA
AT8 all E^HA
S199A^HA
S202A^HA
T205A^HA
AT8 all A^HA
d
AEF of AT8-mutant Tau^HA (a.u.)
Expression intensity (normalized per replicate)
T205E-Tau^HA (*)
S199E^HA (***)
S202E^HA
AT8 all E^HA (*)
e
AEF of AT8-mutant Tau^HA (a.u.)
Expression intensity (normalized per replicate)
S202A^HA
AT8 all A^HA (**)
S199A^HA
T205A^HA
f
AEF of MAP2 (a.u., norm. to 0N3R^HA)
ns
g
DEF of MAP2 (a.u., norm. to 0N3R^HA)
ns
h
NEF of MAP2 (a.u., norm. to 0N3R^HA)
ns

Suppl.Fig. 5

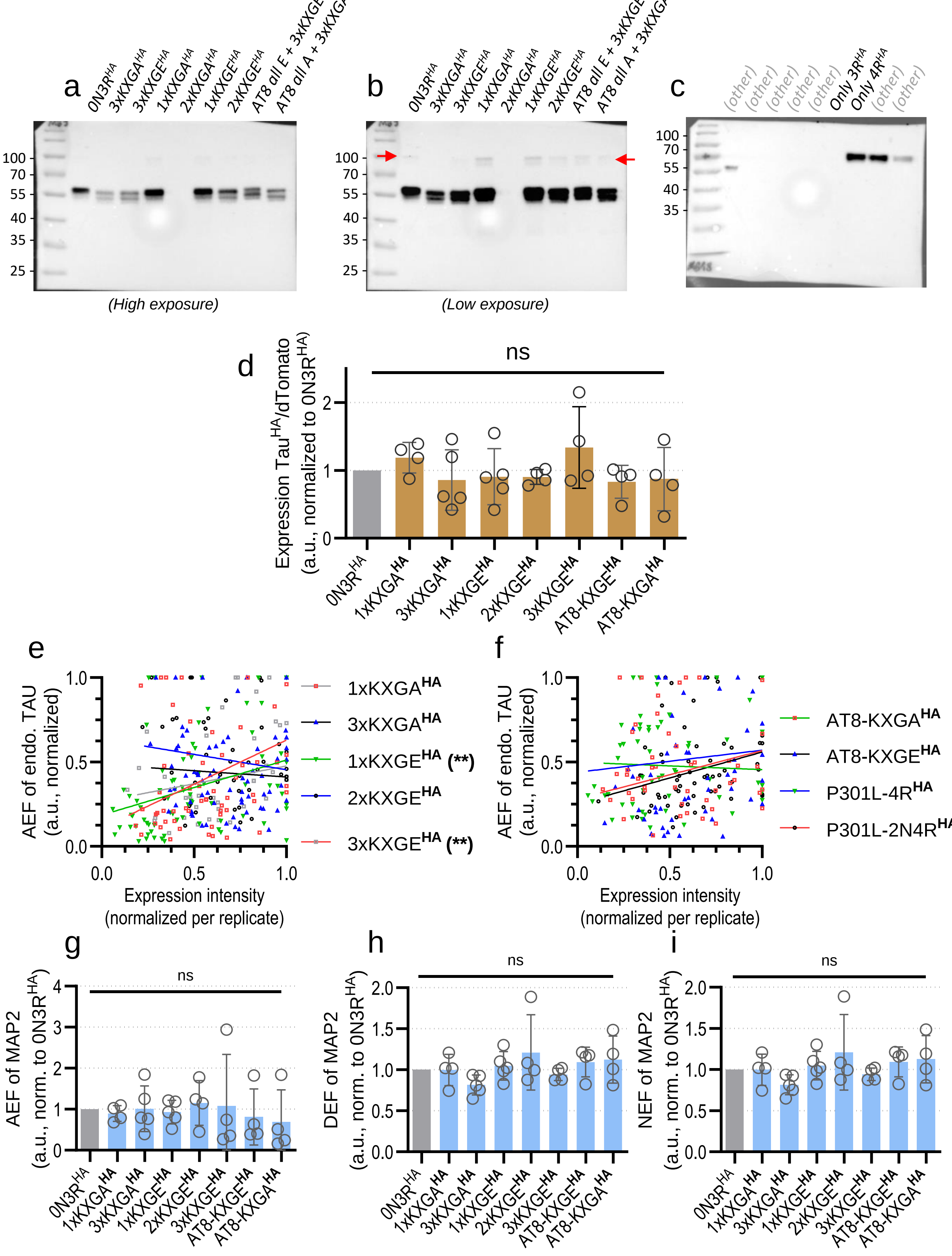
a
0N3R^HA 3xKXGA^HA 3xKXGE^HA 1xKXGA^HA 2xKXGA^HA 1xKXGE^HA 2xKXGE^HA AT8 all E + 3xKXGE^HA AT8 all A + 3xKXGA^HA
(High exposure)
b
(Low exposure)
c
(other) Only 3R^HA Only 4R^HA
d
ns
Expression Tau^HA/dTomato (a.u., normalized to 0N3R^HA)
0N3R^HA 1xKXGA^HA 3xKXGA^HA 1xKXGE^HA 2xKXGE^HA 3xKXGE^HA AT8-KXGE^HA AT8-KXGA^HA
e
AEF of endo. TAU (a.u., normalized)
Expression intensity (normalized per replicate)
1xKXGA^HA
3xKXGA^HA
1xKXGE^HA (**)
2xKXGE^HA
3xKXGE^HA (**)
f
AT8-KXGA^HA
AT8-KXGE^HA
P301L-4R^HA
P301L-2N4R^HA
g
AEF of MAP2 (a.u., norm. to 0N3R^HA)
h
DEF of MAP2 (a.u., norm. to 0N3R^HA)
i
NEF of MAP2 (a.u., norm. to 0N3R^HA)

Suppl.Fig. 6

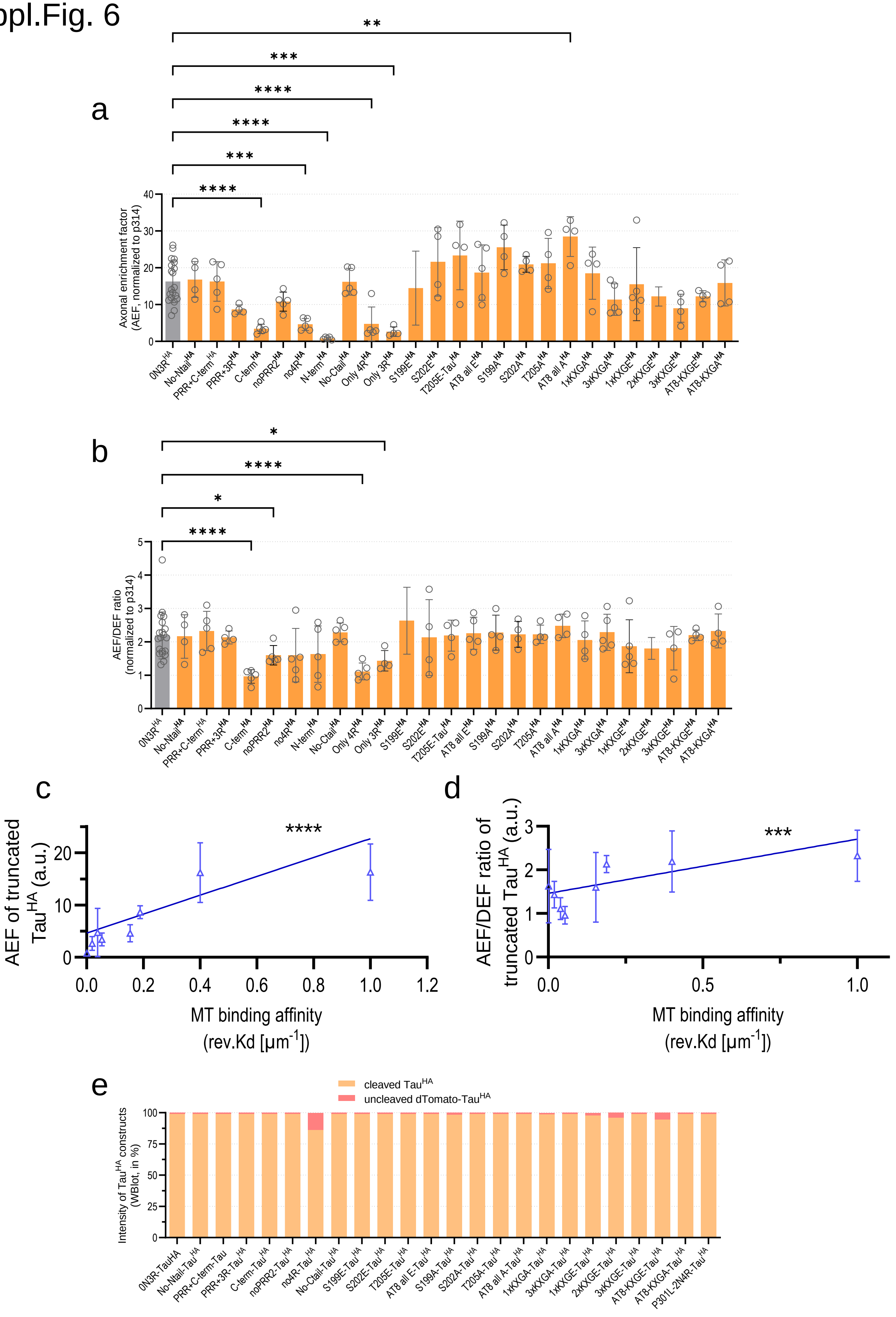

a
Axonal enrichment factor (AEF, normalized to p314)
**
***
****
****
***
****
0N3R^HA No-Ntail^HA PRR+C-term^HA PRR+3R^HA C-term^HA noPRR2^HA no4R^HA N-term^HA No-Ctail^HA Only 4R^HA Only 3R^HA S199E^HA S202E^HA T205E-Tau^HA AT8 all E^HA S199A^HA S202A^HA T205A^HA AT8 all A^HA 1xKXGA^HA 3xKXGA^HA 1xKXGE^HA 2xKXGE^HA 3xKXGE^HA AT8-KXGE^HA AT8-KXGA^HA
b
AEF/DEF ratio (normalized to p314)
*
****
*
****
0N3R^HA No-Ntail^HA PRR+C-term^HA PRR+3R^HA C-term^HA noPRR2^HA no4R^HA N-term^HA No-Ctail^HA Only 4R^HA Only 3R^HA S199E^HA S202E^HA T205E-Tau^HA AT8 all E^HA S199A^HA S202A^HA T205A^HA AT8 all A^HA 1xKXGA^HA 3xKXGA^HA 1xKXGE^HA 2xKXGE^HA 3xKXGE^HA AT8-KXGE^HA AT8-KXGA^HA
c
AEF of truncated Tau^HA (a.u.)
****
MT binding affinity (rev.Kd [µm^-1])
d
AEF/DEF ratio of truncated Tau^HA (a.u.)
***
MT binding affinity (rev.Kd [µm^-1])
e
cleaved Tau^HA
uncleaved dTomato-Tau^HA
Intensity of Tau^HA constructs (WBlot, in %)
0N3R-TauHA No-Ntail-Tau^HA PRR+C-term-Tau PRR+3R-Tau^HA C-term-Tau^HA noPRR2-Tau^HA no4R-Tau^HA No-Ctail-Tau^HA S199E-Tau^HA S202E-Tau^HA T205E-Tau^HA AT8 all E-Tau^HA S199A-Tau^HA S202A-Tau^HA T205A-Tau^HA AT8 all A-Tau^HA 1xKXGA-Tau^HA 3xKXGA-Tau^HA 1xKXGE-Tau^HA 2xKXGE-Tau^HA 3xKXGE-Tau^HA AT8-KXGE-Tau^HA AT8-KXGA-Tau^HA P301L-2N4R-Tau^HA

Suppl.Fig. 7

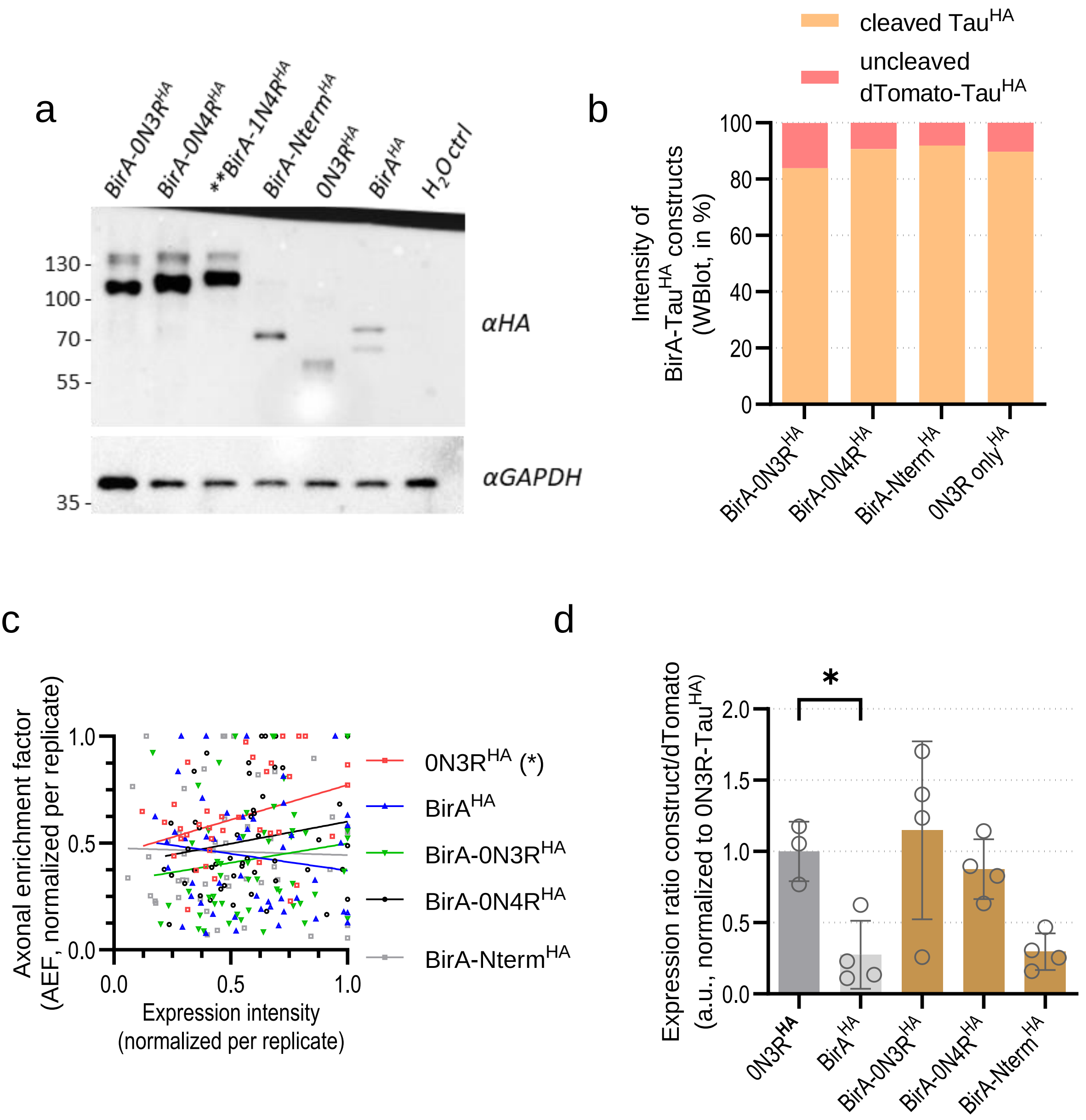
a
BirA-0N3R^HA
BirA-0N4R^HA
**BirA-1N4R^HA
BirA-Nterm^HA
0N3R^HA
BirA^HA
H2O ctrl
130
100
70
55
35
αHA
αGAPDH
b
cleaved Tau^HA
uncleaved dTomato-Tau^HA
Intensity of BirA-Tau^HA constructs (WBlot, in %)
100
80
60
40
20
0
BirA-0N3R^HA
BirA-0N4R^HA
BirA-Nterm^HA
0N3R only^HA
c
Axonal enrichment factor (AEF, normalized per replicate)
1.0
0.5
0.0
Expression intensity (normalized per replicate)
0N3R^HA (*)
BirA^HA
BirA-0N3R^HA
BirA-0N4R^HA
BirA-Nterm^HA
d
*
Expression ratio construct/dTomato (a.u., normalized to 0N3R-Tau^HA)
2.0
1.5
1.0
0.5
0.0
0N3R^HA
BirA^HA
BirA-0N3R^HA
BirA-0N4R^HA
BirA-Nterm^HA

### Suppl.Fig. 1: Distribution of endogenous Tau and MAP2 in differentiating Ngn2-transgenic WTC11 hiPSC-derived neurons.

(a) Differentiation and transduction protocol for *MAPT*-KO iPSC-neurons, based on previously described procedures (Buchholz 2024, Wang 2017, recent paper Li Gan with doxy depletion). 6-weeks-old *MAPT*-KO iPSC-neurons were transduced with lentiviral particles containing the doxycycline-inducible expression vector pUltra-dox for 24 h and kept for 12-13 days in doxycycline-containing medium before harvesting or fixation. (b) Dendritic enrichment (AEF) of endogenous Tau and MAP2 during differentiation normalized to dTomato. Quantification was done for three to five independent experiments (coloured dots) with 8-21 cells per experiment. The coloured dots indicate the arithmetic mean, the error bars show SD. An ordinary two-way ANOVA with Tukey's correction for multiple comparisons was performed to determine significance levels between different weeks and between two DEF values of the same week. Significance levels: * $p < 0.05$, ** $p < 0.01$, ns: $p \geq 0.05$. (c) Nuclear enrichment (NEF) of endogenous Tau and MAP2 during differentiation normalized to dTomato. Quantification was done for three to five independent experiments (coloured dots) with 8-21 cells per experiment. The coloured dots indicate the arithmetic mean, the error bars show SD. An ordinary two-way ANOVA with Tukey's correction for multiple comparisons was performed to determine significance levels between different weeks and between two NEF values of the same week. Significance levels: * $p < 0.05$, ** $p < 0.01$, ns: $p \geq 0.05$.

### Suppl.Fig. 2: Enrichment of transduced 0N3R-Tau$^{HA}$ and non-normalized AEF values of Tau$^{HA}$ constructs.

(a) Tau protein domain structure. The alternatively spliced N-terminal inserts (N1, N2, brown) and C-terminal repeat domains (R1-R4, blue), the Proline-rich domain (subdomains P-rich1, P-rich2, green), and the pseudo-repeat domain (R', grey) are highlighted. The small black numbers indicate the first amino acid of the respective domain, the grey numbers give the corresponding exons of the Tau-encoding *MAPT* gene. The two outlined boxes show motifs (left: AT8 motif, right: KXGS motif) harbouring known highly phosphorylated amino acids residues (bold & underlined). The domain sizes reflect their actual proportion of the whole protein. N = N-terminal insert, R = C-terminal repeat. (b) Dendritic enrichment (DEF) of 0N3R-Tau$^{HA}$ in *MAPT*-KO iPSC-neurons compared to DEF of endogenous Tau in in 6-weeks-old WT iPSC-neurons. (c) Nuclear enrichment (NEF) of 0N3R-Tau$^{HA}$ in *MAPT*-KO iPSC-neurons compared to NEF of endogenous Tau in in 6-weeks-old WT iPSC-neurons. (d) Axonal enrichment (AEF) of endogenous MAP2 in *MAPT*-KO iPSC-neurons expressing 0N3R-Tau$^{HA}$ compared to *MAPT*-KO iPSC-neurons without Tau expression and *MAPT*-WT iPSC-neurons. (e) DEF of endogenous MAP2 in *MAPT*-KO iPSC-neurons expressing 0N3R-Tau$^{HA}$ compared to DEF of *MAPT*-KO iPSC-neurons without Tau expression and *MAPT*-WT iPSC-neurons. (f) NEF of endogenous MAP2 in *MAPT*-KO iPSC-neurons expressing 0N3R-Tau$^{HA}$ compared to NEF of *MAPT*-KO iPSC-neurons without Tau expression and *MAPT*-WT iPSC-neurons. Quantification for (b-f) was done for four to 20 independent experiments (black dots) with 9-21 cells per experiment. The bars indicate the arithmetic mean, error bars show SD. An unpaired t-test (b,c) or ordinary one-way ANOVA with Tukey's correction for multiple comparisons (d-f) was performed to determine significance levels between both groups. Significance levels: ns: * $p < 0.05$, ** $p < 0.01$, ns: $p \geq 0.05$.ns. (g) Correlation of expression levels and axonal enrichment of 0N3R-Tau$^{HA}$ in KO iPSC-neurons. Each data point represents a single neuron. Neurons from all experiments (total of 20) were normalized to the neurons with the maximal expression and maximal enrichment from the respective experiment. Simple linear regression analysis was done for all neurons and indicated a significant positive correlation between the expression level and the axonal enrichment. Significance level: $p < 0.05$, **.

### Suppl.Fig. 3: Characterization of truncated Tau$^{HA}$ construct expression in *MAPT*-KO iPSC-neurons.

(a-c) Unmodified western blots from *MAPT*-KO iPSC-neurons expressing truncated Tau$^{HA}$ constructs, as shown in Fig. 2b. Signal detection was done with anti-HA antibody. Green arrows in (a) highlight Tau$^{HA}$ construct bands after dTomato cleavage, red arrows indicate uncleaved fusion proteins. (d) Protein levels of 0N3R-Tau$^{HA}$ and truncated Tau$^{HA}$ constructs relative to dTomato in *MAPT*-KO iPSC-neurons after 12-13 days of expression, normalized to 0N3R-Tau$^{HA}$. Quantification was done for four to five independent experiments (coloured dots) with 7-16 cells per experiment. The bars indicate the arithmetic mean, error bars show SD. A mixed-effects model with Dunnett's correction for multiple comparisons was performed to determine significance levels between groups. Non-normalized ratios for 0N3R-Tau$^{HA}$ and Tau$^{HA}$ constructs were used for statistical analysis. Significance levels: * $p < 0.05$, ns: $p \geq 0.05$. (e,f) Correlation of expression levels and axonal enrichment of truncated Tau$^{HA}$ constructs in *MAPT*-KO iPSC-neurons. Each data point represents a single neuron (n = 319). Neurons from all experiments (total of 20) were normalized to the neurons with the maximal expression and maximal enrichment from the respective experiment. Simple linear regression analysis was done for all neurons. The slope was significantly different from zero for noPRR2-Tau$^{HA}$ (e, positive correlation, p = 0.0044) and only4R-Tau$^{HA}$ (f, positive correlation, p = 0.0269), and not significantly different for all other truncated Tau$^{HA}$ constructs. (g) Axonal enrichment (AEF) of endogenous MAP2 in *MAPT*-KO iPSC-neurons after 12-13 days of truncated Tau$^{HA}$ construct expression, normalized to AEF of endogenous MAP2 in *MAPT*-KO iPSC-neurons expressing 0N3R-Tau$^{HA}$. (h) Dendritic enrichment (DEF) of endogenous MAP2 in *MAPT*-KO iPSC-neurons after 12-13 days of truncated Tau$^{HA}$ construct expression, normalized to DEF of endogenous MAP2 in *MAPT*-KO iPSC-neurons expressing 0N3R-Tau$^{HA}$. (i) Nuclear enrichment (NEF) of endogenous MAP2 in *MAPT*-KO iPSC-neurons after 12-13 days of truncated Tau$^{HA}$ construct expression, normalized to NEF of endogenous MAP2 in *MAPT*-KO iPSC-neurons expressing 0N3R-Tau$^{HA}$. Quantification for (f-h) was done for four to five independent experiments (black dots) with 8-19 cells per experiment. The bars indicate the arithmetic mean, error bars show SD. A mixed-effects model with Dunnett's correction for multiple comparisons was performed to determine significance levels between 0N3R-Tau$^{HA}$ and all Tau$^{HA}$ constructs. Non-normalized ratios of endogenous MAP2 were used for the analysis. Significance level: ns: $p \geq 0.05$.

### Suppl.Fig. 4: Characterization of AT8-mutant Tau$^{HA}$ construct expression in *MAPT*-KO iPSC-neurons.

(a,b) Unmodified western blots from *MAPT*-KO iPSC-neurons expressing AT8-mutant Tau$^{HA}$ constructs, as shown in Fig. 3b. Signal detection was done with anti-HA antibody. Red arrows indicate uncleaved dTomato fusion proteins barely visible in some lanes (a). (c) Protein levels of 0N3R-Tau$^{HA}$ and AT8-mutant Tau$^{HA}$ constructs relative to dTomato in *MAPT*-KO iPSC-neurons after 12-13 days of expression, normalized to 0N3R-Tau$^{HA}$. Quantification was done for four to five independent experiments (coloured dots) with 9-17 cells per experiment. The bars indicate the arithmetic mean, error bars show SD. A mixed-effects model with Dunnett's correction for multiple comparisons was performed to determine significance levels between groups. Non-normalized ratios for 0N3R-Tau$^{HA}$ and Tau$^{HA}$ constructs were used for statistical analysis. Significance levels: * $p < 0.05$, ns: $p \geq 0.05$. (d,e) Correlation of expression levels and axonal enrichment of AT8-mutant Tau$^{HA}$ constructs in *MAPT*-KO iPSC-neurons. Each data point represents a single neuron. Neurons from all experiments (total of 20) were normalized to the neurons with the maximal expression and maximal enrichment from the respective experiment. Simple linear regression analysis was done for all neurons. The slope was significantly different from zero for S199E$^{HA}$ (d, positive correlation, p = 0.0002), T205E$^{HA}$ (d, positive correlation, p = 0.0412), AT8 allE$^{HA}$ (d, positive correlation, p = 0.0431), and AT8 allA$^{HA}$ (e, positive correlation, p = 0.0023), and not significantly different for all other AT8-mutant Tau$^{HA}$ constructs. (f) Axonal enrichment (AEF) of endogenous MAP2 in *MAPT*-KO iPSC-neurons after 12-13 days of AT8-mutant Tau$^{HA}$ construct expression, normalized to AEF of endogenous MAP2 in *MAPT*-KO iPSC-neurons expressing 0N3R-Tau$^{HA}$. (g) Dendritic enrichment (DEF) of endogenous MAP2 in *MAPT*-KO iPSC-neurons after 12-13 days of AT8-mutant Tau$^{HA}$ construct expression, normalized to DEF of endogenous MAP2 in *MAPT*-KO iPSC-neurons expressing 0N3R-Tau$^{HA}$. (h) Nuclear enrichment (NEF) of endogenous MAP2 in *MAPT*-KO iPSC-neurons after 12-13 days of AT8-mutant Tau$^{HA}$ construct expression, normalized to NEF of endogenous MAP2 in *MAPT*-KO iPSC-neurons expressing 0N3R-Tau$^{HA}$. Quantification for (f-h) was done for four to five independent experiments (black dots) with 9-17 cells per experiment. The bars indicate the arithmetic mean, error bars show SD. A mixed-effects model with Dunnett's correction for multiple comparisons was performed to determine significance levels between 0N3R-Tau$^{HA}$ and all Tau$^{HA}$ constructs. Non-normalized ratios of endogenous MAP2 (h) were used for the analysis. Significance level: ns: $p \geq 0.05$.

### Suppl.Fig. 5: Characterization of KXGS- and double-mutant Tau$^{HA}$ construct expression in *MAPT*-KO iPSC-neurons.

(a-c) Unmodified western blots from *MAPT*-KO iPSC-neurons expressing KXGS- and double-mutant Tau$^{HA}$ constructs, as shown in Fig. 4b. Signal detection was done with anti-HA antibody. Red arrows indicate uncleaved dTomato fusion proteins barely visible in some lanes (b). (d) Protein levels of 0N3R-Tau$^{HA}$ and KXGS- and double-mutant Tau$^{HA}$ constructs relative to dTomato in *MAPT*-KO iPSC-neurons after 12-13 days of expression, normalized to 0N3R-Tau$^{HA}$. Quantification was done for four to five independent experiments (coloured dots) with 9-17 cells per experiment. The bars indicate the arithmetic mean, error bars show SD. A mixed-effects model with Dunnett's correction for multiple comparisons was performed to determine significance levels between groups. Non-normalized ratios for 0N3R-Tau$^{HA}$ and Tau$^{HA}$ constructs were used for statistical analysis. Significance levels: * $p < 0.05$, ns: $p \geq 0.05$. (e,f) Correlation of expression levels and axonal enrichment of KXGS- and double-mutant Tau$^{HA}$ constructs in *MAPT*-KO iPSC-neurons. Each data point represents a single neuron. Neurons from all experiments (total of 20) were normalized to the neurons with the maximal expression and maximal enrichment from the respective experiment. Simple linear regression analysis was done for all neurons. The slope was significantly different from zero for 1xKXGE$^{HA}$ (e, positive correlation, $p = 0.0044$), 3xKXGE$^{HA}$ (e, positive correlation, $p = 0.0034$), and not significantly different for all other KXGS- and double-mutant Tau$^{HA}$ constructs. (f) Axonal enrichment (AEF) of endogenous MAP2 in *MAPT*-KO iPSC-neurons after 12-13 days of KXGS- and double-mutant Tau$^{HA}$ construct expression, normalized to AEF of endogenous MAP2 in *MAPT*-KO iPSC-neurons expressing 0N3R-Tau$^{HA}$. (g) Dendritic enrichment (DEF) of endogenous MAP2 in *MAPT*-KO iPSC-neurons after 12-13 days of KXGS- and double -mutant Tau$^{HA}$ construct expression, normalized to DEF of endogenous MAP2 in *MAPT*-KO iPSC-neurons expressing 0N3R-Tau$^{HA}$. (h) Nuclear enrichment (NEF) of endogenous MAP2 in *MAPT*-KO iPSC-neurons after 12-13 days of KXGS- and double-mutant Tau$^{HA}$ construct expression, normalized to NEF of endogenous MAP2 in *MAPT*-KO iPSC-neurons expressing 0N3R-Tau$^{HA}$. Quantification for (f-h) was done for four to five independent experiments (black dots) with 8-19 cells per experiment. The bars indicate the arithmetic mean, error bars show SD. A mixed-effects model with Dunnett's correction for multiple comparisons was performed to determine significance levels between 0N3R-Tau$^{HA}$ and all Tau$^{HA}$ constructs. Non-normalized ratios of endogenous MAP2 were used for the analysis. Significance level: ns: $p \geq 0.05$.

### Suppl.Fig. 6: Non-normalized AEF values, AEF/DEF ratio, AEF/MT binding ratio, and cleavage efficiency of all Tau$^{HA}$ constructs.

(a) Non-normalized axonal enrichment (AEF) of 0N3R-Tau$^{HA}$ (grey) and all Tau$^{HA}$ constructs (orange). Quantification was done for four to five independent experiments (black dots) with 8-19 cells per experiment. The bars indicate the arithmetic mean, error bars show SD. A mixed-effects model with Dunnett's correction for multiple comparisons was performed to determine significance levels between 0N3R-Tau$^{HA}$ and all Tau$^{HA}$ constructs. Significance levels: ** $p < 0.01$, *** $p < 0.001$, **** $p < 0.0001$, ns: $p \geq 0.05$. (b) Non-normalized ratio of axonal to dendritic enrichment (AEF/DEF) of 0N3R-Tau$^{HA}$ (grey) and all Tau$^{HA}$ constructs (orange). Quantification was done for four to five independent experiments (black dots) with 8-19 cells per experiment. The bars indicate the arithmetic mean, error bars show SD. A mixed-effects model with Dunnett's correction for multiple comparisons was performed to determine significance levels between 0N3R-Tau$^{HA}$ and all Tau$^{HA}$ constructs. Significance levels: * $p < 0.05$, **** $p < 0.0001$, ns: $p \geq 0.05$. (c) AEF of all truncated Tau$^{HA}$ constructs reversely correlated to their microtubule binding affinity (Gustke et al. 1994). A simple linear regression analysis was performed. The slope was significantly different from zero (positive correlation, <p = 0.0001). (d) AEF/DEF ratio of all truncated Tau$^{HA}$ constructs reversely correlated to their microtubule binding affinity (Gustke et al. 1994). A simple linear regression analysis was performed. The slope was significantly different from zero (positive correlation, $p = 0.0003$). (c) Proportions of Tau$^{HA}$ constructs (orange) and uncleaved dTomato-Tau$^{HA}$ fusion proteins (red), quantified from Western blots (Figs. 2b,3b,4b). Western blot with different exposure times were used for quantification to avoid saturation but ensure optimal detection of fainter bands. Except for no4R$^{HA}$, which had a proportion of uncleaved protein of ~14 %, all constructs were cleaved from dTomato with at least 95 % efficiency. Quantification was done for one replicate per Tau$^{HA}$ construct.

### Suppl.Fig. 7: Characterization of BirA-Tau$^{HA}$ construct expression in *MAPT*-KO iPSC-neurons.

(a) Western blot of *MAPT*-KO iPSC-neurons transduced with BirA-Tau$^{HA}$ constructs. GAPDH and was used as loading control. Distinct bands are visible for the three isoform fusion constructs and BirA-Nterm-Tau$^{HA}$, with weak high-molecular signals deriving from uncleaved Tau$^{HA}$-P2A-dTomato fusion proteins (see panel b). BirA-1N4R$^{HA}$ was included for Western blot analysis but not used elsewhere in this study. (b) Proportions of cleaved BirA-Tau$^{HA}$ constructs (orange parts) and uncleaved dTomato-Tau$^{HA}$ fusion proteins (red parts), quantified from blotted protein lysates (Figs. 2b,3b,4b). Western blot with different exposure times were used for quantification to avoid saturation but ensure optimal detection of fainter bands. The cleavage efficiency ranged from 84 % (BirA-0N3R$^{HA}$) to 92 % (BirA-Nterm$^{HA}$). (c) Correlation of expression levels and axonal enrichment of BirA-Tau$^{HA}$ constructs in *MAPT*-KO iPSC-neurons. Each data point represents a single neuron. Neurons from all experiments (total of 20) were normalized to the neurons with the maximal expression and maximal enrichment from the respective experiment. Simple linear regression analysis was done for all neurons. The slope was significantly different from zero for 0N3R-Tau$^{HA}$ (positive correlation, $p = 0.0472$), and not significantly different for all other BirA-Tau$^{HA}$ constructs. (d) Protein levels of 0N3R-Tau$^{HA}$ and BirA-Tau$^{HA}$ constructs relative to dTomato in *MAPT*-KO iPSC-neurons after 12-13 days of expression, normalized to 0N3R-Tau$^{HA}$. Quantification was done for three to four independent experiments (coloured dots) with 9-17 cells per experiment. The bars indicate the arithmetic mean, error bars show SD. An ordinary one-way ANOVA with Dunnett's correction for multiple comparisons was performed to determine significance levels between groups. Non-normalized ratios for 0N3R-Tau$^{HA}$ and BirA-Tau$^{HA}$ constructs were used for statistical analysis. Significance levels: ns: $p \geq 0.05$.
