## Supplementary material for "Axonal Tau sorting depends on the PRR2 domain and 0N4R-specific interactions hint at distinct roles of Tau isoforms in synaptic plasticity": Supplee

#### MACRO #1: File renamer and saver ###

// Set the directory path where you want to save the ROI files
path = "U:/winfiles/Desktop/Test";
// Define the new names for the ROIs
roiNames = newArray("axon", "axon background", "soma", "soma background");


//script start

// Open each CZI file in the folder
list = getFileList(path + "/Images");
Array.sort(list);

//define variable for the images
for (j =0; j < list.length; j++) {
	//open the images
	if(endsWith(list[j], ".czi")) {
    	run("Bio-Formats Importer", "open=[" + path + "/Images/" + list[j] + "]");
		
		run("Split Channels");

		// Construct the full path to the ROI file by joining the directory path and the filename
		roiFile = "/ROI/" + File.getNameWithoutExtension(list[j]) + ".zip";
		
		 // Wait for user confirmation before proceeding with ROIs
    	waitForUser("Press OK to save and rename ROIs");
		
		
		// Loop through each ROI and rename it
		for (i=0; i<roiManager("count"); i++) {
  		roiManager("select", i);
  		roiManager("rename", roiNames[i]);
  		roiManager("Remove Channel Info");
		roiManager("Remove Slice Info");
		roiManager("Remove Frame Info");
  		if (i == roiNames.length-1) break; // exit loop when all new names have been used
		}


	// Save the ROI set with the name of the open image
	roiManager("save", path + roiFile);

	//reset all
	close("*");
	roiManager("reset")
	}
}


### MACRO #2: Automated ROI measurement ###

//i-loop iterates over the ROIs, while the inner j-loop opens all images (long script), plan to extract results as single columns


// Set the path to the folder containing the CZI files and ROI collections
//The Folder should contain three subfolders: 
//"Images" (containing only .czi files), 
//"ROI" (containing only .zip files with the name corresponding to the .czi files,
//"Results" - This script overwrites all files with the same name as the result files.
path = "U:/winfiles/Desktop/BlindRandomFiles2023_05_05_T_13_06_16_120/01"

//Define the names of the ROIs within your ROI set, these will be appended to your results files
roiNames = newArray("soma", "soma_background", "nucleus");


//script start

//loop through all the channels with c
for (c=1; c<5;c++){
	SetChannel=c;
	
	//define the image-identifier 
	channel = "C" + SetChannel + "-";
	
	// Create a new table for the results
	Table.create("Results");

	// Open each CZI file in the folder
	list = getFileList(path + "/Images");

	// Sort the list of files alphabetically
	Array.sort(list);

	//define variable for which ROI to use
	for (i = 0; i < 3; i++) {
    
		//define variable for the images
		for (j =0; j < list.length; j++) {
			//open the images
			if(endsWith(list[j], ".czi")) {
    			run("Bio-Formats Importer", "open=[" + path + "/Images/" + list[j] + "]");

				// Split the channels
				run("Split Channels");

    			selectWindow(channel + path + "/Images/" + list[j]);
    		
        		// Find the corresponding ROI collection by name
				roiFile = File.getNameWithoutExtension(list[j]) + ".zip";
	    		roiSetPath = path + "/ROI/" + roiFile;
        		roiManager("Open", roiSetPath);
        		//open the i-th ROI
        		roiManager("Select", i)
        		// Measure the mean intensity
        		run("Measure","mean");
				
       			// Close the CZI file and ROI collection
        		close("*");
        		roiManager("reset")
			}
		}

		//extract image id
		nRows=nResults;
		image_id=newArray(nRows);
		for (id=0; id<list.length; id++) {
			image_id[id]=list[id];
		}
	
		//create and empty array for the results
		nRows=nResults;
		extract = newArray(nRows);
		//loop through the results table
		for (r=0; r<nRows; r++){
			extract[r]=getResult("Mean", r);
		}
	
		Table.create(channel+"Result");
		Table.setColumn(roiNames[i], extract);
		Table.setColumn("image_id", image_id)
		run("Clear Results");
		// Save the results table to a CSV file
		Table.save(path + "/Results/" + channel + "result_" + roiNames[i] + ".csv");
	
		image_id=newArray(0);
	}
}
